# A multichromatic UV-RGB optogenetic toolbox for control of gene expression in *Pseudomonas putida*

**DOI:** 10.64898/2026.05.27.727800

**Authors:** Seung-Hyun Paik, Gi-Hyun Paik, Luzie Kruse, Tobias Horbach, Fabienne Hilgers, Andrea Weiler, Matthias Pesch, Lennart Witting, Michelle Bund, Dietrich Kohlheyer, Thomas Drepper

## Abstract

Optogenetics uses light to provide precise, reversible, and non-invasive control over bacterial functions including gene expression with high spatiotemporal resolution. Although many optogenetic systems have been developed for *Escherichia coli*, only a limited number is available for other prokaryotes, such as pseudomonads. Here, we establish a toolbox of genomically integrated optogenetic gene cassettes for light-responsive regulation of target gene expression in *Pseudomonas putida* with UV-A, blue, green, and red light. Using transposon Tn7-mediated chromosomal integration, we implemented four optogenetic systems: the photocaged IPTG/P_tac_-LacI system, the LOV-based Dusk switch, the cyanobacteriochrome system CcaS/R, and different bacteriophytochrome-based REDusk variants. Benchmarking with the mCherry reporter demonstrated high dynamic ranges of up to ∼270-fold, low basal expression, and largely homogeneous population responses in *P. putida*. Spatial illumination further enabled patterned single- and dual-color gene expression. As a proof of concept, we applied the toolbox for light-controlled regulation of pyoverdine (PVD) biosynthesis in *P. putida*. The expression of the alternative sigma factor PfrI, which upregulates the production of the siderophore during iron-limitation, was placed under optogenetic control in a Δ*pfrI* background. The red-light responsive switches resulted in the strongest induction of PVD synthesis and enabled spatial control of siderophore-mediated microbial interactions. To demonstrate transferability, light-dependent pyoverdine production was further established in the human pathogen *Pseudomonas aeruginosa* PAO1. Together, this optogenetic plug-and-play toolbox enables non-invasive, spatiotemporal reprogramming of gene expression and cellular processes in pseudomonads and expands the available optogenetic repertoire beyond established model organisms.

## Introduction

In bacterial optogenetics, the use of light-responsive switches has revolutionized the field of synthetic biology by providing precise, non-invasive control over cellular functions including gene expression. Implementing synthetic, light-controlled regulatory circuits thereby enables the non-invasive reprogramming of almost any bacterial function and cellular behavior with high temporal and spatial resolution^1^. Optogenetic switches are typically based on engineered photoreceptors and optochemical switches that respond specifically to light excitation at a defined wavelength. Different photoreceptor classes, such as light-oxygen-voltage (LOV) domains ^2^ (blue light, flavin mononucleotide as chromophore), bacteriophytochromes^3,4^ (red/far-red light, biliverdin as chromophore), cyanobacteriochromes^5^ (green/red light, e.g., phycocyanobilin as chromophore) or caged-inducers^6,7^ (UV light, e.g., 6-nitropiperonyl, NP, as photocage) provide a wide spectral range for regulation. Engineered transcriptional regulators or two-component systems (TCS) in combination with their corresponding promoters can therefore be used as dynamic, reversible and modular switches for light-triggered gene expression. Well established examples of optogenetic switches in bacteria include the CcaS/R system ^5,8,9^, which responds to green and red light, LOV-based repressor/activator system such as the dusk/dawn switches ^10^or the photocaged isopropylβ-D-1-thiogalactopyranoside (cIPTG)^11^. While primarily *E. coli* has been used as a model to develop and optimize these systems, applications in other bacteria remain limited. For example, for the metabolically versatile strain *Pseudomonas putida* KT2440, so far only a few optogenetic switches have been used, such as the TCS CcaS/R^12–14^ and caged inducers such as NP-cIPTG.^15^ For CcaS/R, adapting and optimizing the cyanobacterial TCS enabled green-light-responsive control of heterologous gene expression in *P. putida*, which its applicability was further demonstrated by light-regulated biofilm formation^14^. In addition, photocaged IPTG derivatives were shown to allow for UV-A mediated activation of LacI-controlled gene expression in *P. putida*, with increased inducer solubility leading to higher expression levels^15^. Expanding the optogenetic toolbox for light-regulated gene expression in *P. putida* is particularly interesting, as this organism is environmentally robust making it an excellent host for bioproduction, bioremediation and synthetic biology.^16–19^

In this study, we present a toolbox for light-controlled gene expression in *P. putida*, comprising optochemical and optogenetic switches responsive to UV-A (∼ 365 nm), blue (∼ 450 nm), green (∼ 530 nm) and red (∼ 660 nm) light. These switches enable precise and customizable control of genetic and metabolic activities. To demonstrate and validate the applicability of the new toolbox, we established dynamic, light-dependent control of pyoverdine (PVD) biosynthesis, a siderophore of major ecological and biotechnological relevance.^20–23^ In addition, we transferred the same regulatory strategy to *Pseudomonas aeruginosa,* an opportunistic human pathogen, and achieved light regulated PVD production, highlighting the broad application potential within the genus *Pseudomonas* and its relevance for biomedical applications.^24,25^ By expanding the repertoire of optogenetic tools available for *P. putida,* this work provides a foundation for multifactorial, spatiotemporal programming of microbial functions applicable in metabolic engineering, engineering of microbial consortia and synthetic biology.

## Results and discussion

### Design and structure of the optogenetic toolbox for UV-RGB light-mediated control of gene expression in *P. putida*

To establish a versatile optogenetic toolbox for UV-RGB light-controlled gene expression in *Pseudomonas putida*, we selected different optochemical and optogenetic switches that cover four distinct spectral ranges (Figure 1, Tab. S2): **UV**-A (∼ 365 nm), **R**ed (∼ 650 nm), **G**reen (∼ 530 nm) and **B**lue (∼ 450 nm). For the UV-A responsive switch, we chose NP-cIPTG (irreversible **ON** switch) in combination with a P_tac_-LacI system (Figure 1 A,B **I**,^7,26^). In the presence of the photo-removable protection group 6-nitropiperonyl, IPTG is biologically inactive and LacI represses the P_tac_ promoter. Upon illumination with UV-A light the NP group is cleaved off, releasing the bioactive inducer molecule. Subsequently, IPTG can bind to LacI, leading to the derepression of the P_tac_ promoter that mediates target gene expression (gene of interest, GOI). For the red-light controlled expression systems, we evaluated the bacteriophytochrome-based photoreceptors rREDusk, rDERusk, mREDusk, mDERusk^3,4^. These switches derived from phytochromes and bind billiverdine IXα as a chromophore. rREDusk contains a photosensing module (PSM), derived from *Deinococcus radiodurans*, fused to histidine kinase of *Bradyrhizobium japonicum* (FixL)(Figure 1 A,B **II,**^3,4^). In the absence of light, the kinase domain of the fusion protein autophosphorylates and transfers the phosphate to the response regulator FixJ, which then activates FixK2 promoter-dependent transcription of the GOI. Upon illumination with red light, the system shifts towards a phosphatase-activated state, leading to dephosphorylation of FixJ and repression of P_FixK2_-driven transcription. Recently, the light sensitivity and mode of action could be engineered by integrating the PSM of a phytochrome derived from *Deinococcus maricopensis* and by varying the linker length between the kinase and sensor domain. Through these modifications rDERusk (reversible red **ON** switch, linker modification), mREDusk (reversible red **OFF** switch) with PSM from *D. maricopensis* and mDERusk (reversible red **ON** switch, linker modification) with PSM from *D. maricopensis* were generated^3^. For the green light regulated optogenetic system, we used the CcaS/R based system, originally derived from *Synechocystis sp.* PCC6803 (Figure 1 A,B **III**,^5,9^). CcaS is a sensor-histidine kinase, which binds phycocyanobilin as chromophore. Illumination with green light promotes autophosphorylation of CcaS, transferring the phosphoryl group to the response regulator CcaR, which then binds to the promoter P_cpcG2-172_, leading to GOI transcription. This process is reversible upon illumination with red light (reversible green **ON**/red **OFF** switch). This system was already previously characterized and successfully implemented in *P. putida* ^12–14^. Lastly, for the blue light regulation of GOI expression we chose Dusk (reversible blue **OFF** switch), which is based on an FMN-binding LOV-domain photoreceptor (Fig. 1 A, B **IV**,^10^). Dusk is a fusion protein (YF1) that consists of the LOV domain from the *Bacillus subtilis* YtvA photoreceptor that has been fused to the histidine kinase domain of FixL and was used as a scaffold to generate the REDusk variants. In darkness YF1 phosphorylates its response regulator FixJ thereby activating the FixK2 promoter. Upon illumination this process is reversed analog to the REDusk system. A similar system named Dawn, in which FixJ controls the expression of a C1 repressor, for the inversion of the signal (blue **ON** switch) was already used in *P. aeruginosa* to control the expression of genes involved in biofilm formation^27^. This demonstrates that YF1-based blue-light control is functional in pseudomonads strongly suggesting that similar modules can work in *P. putida.* For the formation of photoreceptor-based optogenetic switches, the corresponding chromophores have to be supplied by *Pseudomonas* (i.e., FMN synthesized by *P. putida* for YF1), or two additional genes must be heterologously expressed (i.e., the heme oxygenase gene *ho1* for biliverdine synthesis and additionally the ferredoxin oxidoreductase gene *pcyA* for phycocyanobiline production, Figure 1A ^4,5,10^).

**Figure 1.**
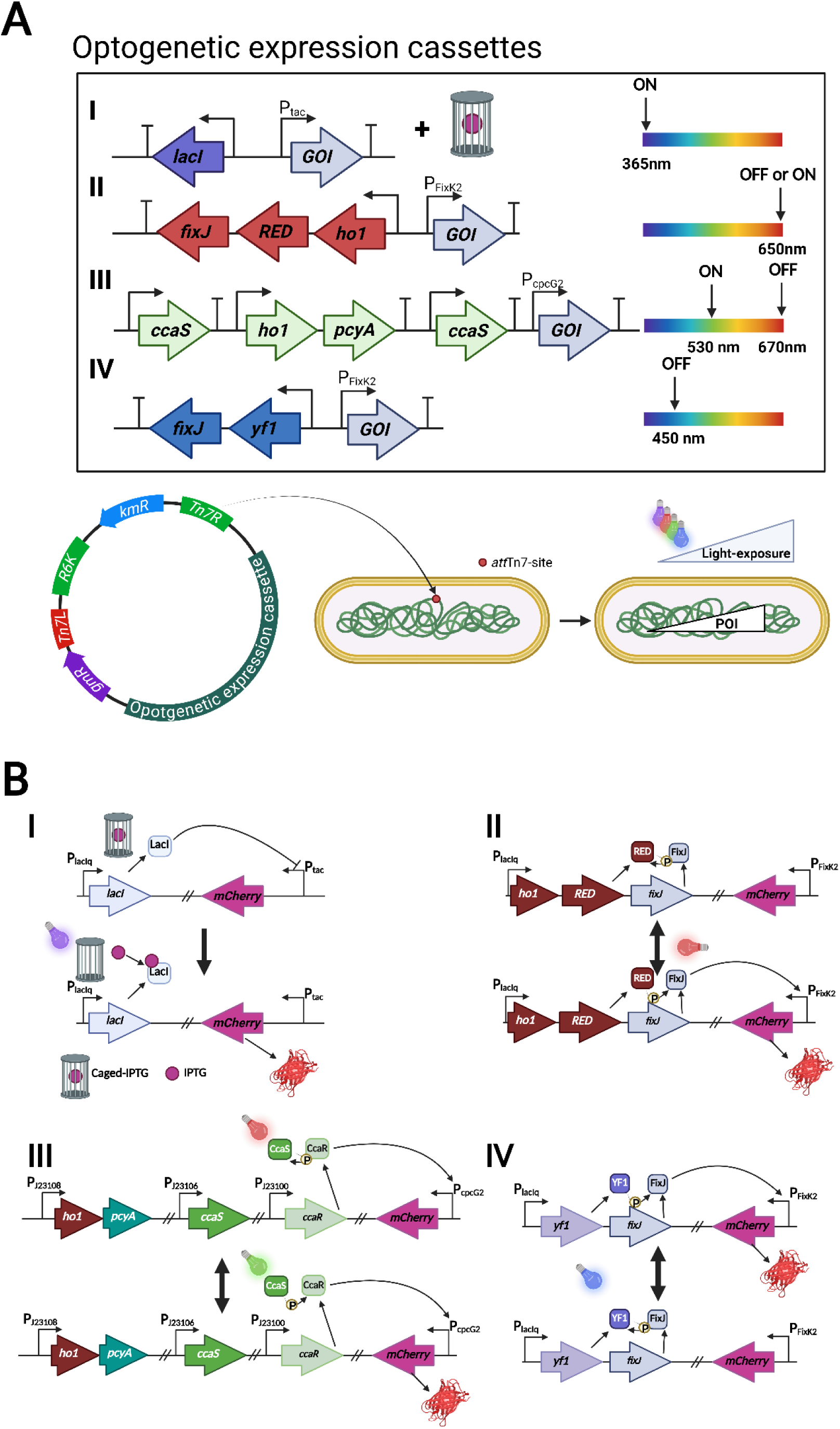
Architecture of gene cassettes encoding UV-RGB-controllable optogenetic switches for site-specific, Tn7-mediated integration into the chromosome of *P. putida*. (A) Structure of the optogenetic cassettes (I: P_tac_-LacI+NP-cIPTG; II: RED-light sensitive systems rREDusk/rDERusk/mREDusk/mDERusk; III: CcaS/R; IV: Dusk) and schematic overview of the genomic integration strategy. Optogenetic modules, designed for light-dependent regulation of gene of interest (GOI) expression, were cloned into a Tn7-based integration vector for stable insertion into the *P. putida* chromosome (Tn7-specific *att*Tn7 site^28^). (B) Overview about the function of the selected optogenetic switches, here controlling the expression of mCherry. (I) P_tac_–LacI + NP-cIPTG (UV-A, ∼365 nm): UV-A light illumination releases IPTG from the caged inducer cIPTG, derepressing LacI-controlled promoters and enabling GOI expression. (II) rDERusk/rREDusk and mDERusk/mREDusk (red, 650 nm): red-light responsive two-component systems (RED/FixJ/P_FixK2_) enables ON (rDERusk/mDERusk) or OFF (rREDusk/mREDusk) regulation of GOI transcription. (III) CcaS/R (green/red, ∼530 nm / 670 nm): A green-light activated, red light repressed two-component system allows dual-color control of gene expression. (IV) Dusk (blue, 450 nm): A blue-light sensitive LOV–histidine kinase (YF1/FixJ) enables control of GOI transcription as an OFF switch

While most of the so far applied optogenetic systems in bacteria are plasmid-encoded^1^, we consciously chose to integrate our expression cassettes into the chromosome of *P. putida*. Chromosomal integration offers several advantages: it ensures stable maintenance of the optogenetic cassette without the need for continuous antibiotic selection, it eliminates copy number related functional variabilities caused by plasmid replication or unequal distribution during cell division, and it reduces metabolic burden, which can be caused by plasmid replication^29–32^. The lack of antibiotic selection is especially important, when studying microbial or plant-microbe interactions^33^, while lowering metabolic burden can be important for the optimization of biotechnological production processes^34^. For genomic integration we employed the Tn7-based transposon system, that mediates site-specific integration at the *att*Tn7-site downstream of the *glmS* gene, a gene locus that is highly conserved in bacteria. This system allows fast and stable single copy insertions into a functionally neutral chromosomal insertion site, therefore minimizing the risk of potentially disrupting the hosts physiology. Furthermore, Tn7-based chromosomal engineering has been widely applied for use in various *Pseudomonas* species and especially *P. putida*, being a favorable tool for directed genomic integration^35,36^. In summary, the toolbox design combines multifactorial light control with stable genomic integration suitable for application in *P. putida* and other pseudomonads.

### Comparative characterization of the UV-RGB light-switches in *P. putida*

To characterize and benchmark the optogenetic switches *in vivo*, the fluorescent reporter gene *mCherry* was placed downstream of the different light-controllable promoters and the resulting expression cassettes were genomically integrated into the *att*Tn7 site of *P. putida* KT2440, generating *P. putida::att*Tn7-P*_tac_*-mCherry (NP), *P. putida::att*Tn7-rREDusk-mCherry (rR), *P. putida::att*Tn7-rDERusk-mCherry (rD), *P. putida::att*Tn7-mREDusk-mCherry (mR), *P. putida::att*Tn7-mDERusk-mCherry (mD), *P. putida::att*Tn7-CcaS/R-mCherry (CcaS/R) and *P. putida::att*Tn7-Dusk-mCherry(Dusk) (Figure 2).

**Figure 2.**
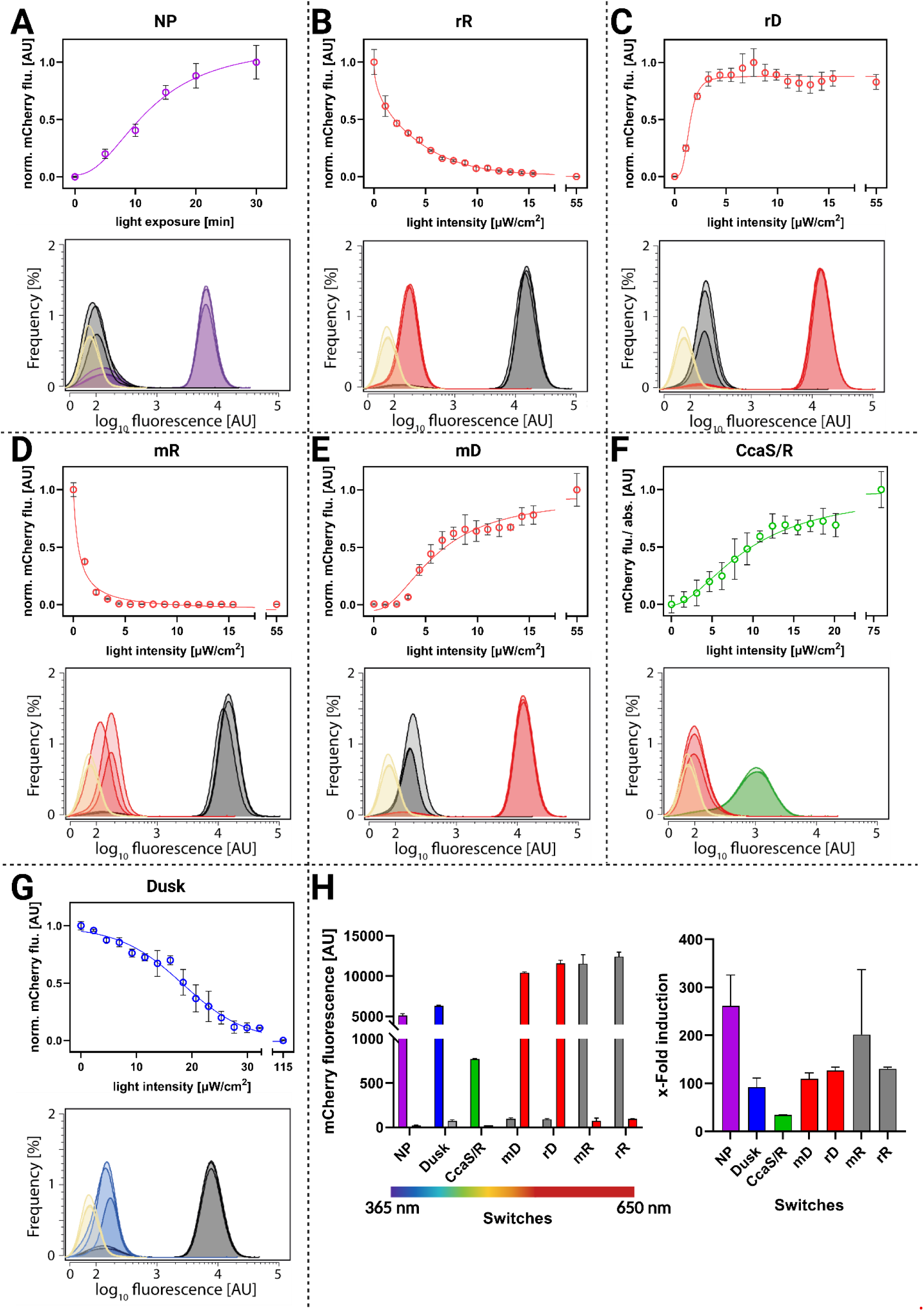
UV-RGB light-controlled gene expression in *Pseudomonas putida*. *P. putida* strains carrying genomically integrated optogenetic expression cassettes were cultivated in 1 mL LB medium (1200rpm, 30°C) under varying illumination conditions and wavelengths using a programmable LED matrix (BioLight, Figure S3, Table S1). The tested systems comprised (A) P_tac_–LacI combined with NP-cIPTG (NP, UV-A responsive, ∼365 nm),(B) the red-light OFF switch rREDusk (rR, 620–625 nm), (C) the red light ON switch rDERusk (rD, 620–625 nm), (D) the red light OFF switch mREDusk(mR, 620–625 nm), (E) the red light ON switch mDERusk(mD, 620–625 nm), (F) the green/red two-component system CcaS/R (ON: ∼522–525 nm and OFF: ∼620–625 nm), and (G) the blue-light OFF switch Dusk (465–467 nm). The *P. putida* NP cultures were individually illuminated after 4 hours for 0-30 mins of UV-A light (365nm ≈ 1mW/cm^2^) for the gradual uncaging of NP-cIPTG. *P. putida* strains harboring the different photoreceptor systems (rR, rD, mR, mD, CasS/R, and Dusk) were cultivated under individually increasing light intensities at the respective wavelengths or in the dark. After 24 hours, mCherry fluorescence (λ_ex_ = 575 nm, λ_em_ = 610 nm) was measured in all *P. putida* cultures. Line plots (top) show mCherry fluorescence normalized to the respective cell densities as a function of light exposure time or intensity. Data represents mean values ± standard deviation (SD) from biological triplicates. Flow cytometry histograms (bottom) display single-cell mCherry fluorescence distributions of 10,000 measured events. Colors correspond to the applied illumination conditions (longest illumination time for NP or highest light intensities): A: purple (NP, UV-A; ON), F: green (CcaS/R; ON), red (CcaS/R; OFF), B-E: red (rR, rD, mR, and mD; ON or OFF), G: blue (Dusk; OFF), black (darkness), and yellow (WT control without optogenetic expression cassettes). All experiments displayed in biological triplicates from three different cultivations. (H) Left: Summary of all tested optogenetic switches including their respective spectral coverage, and the average mCherry fluorescence of 10,000 tracked events under fully induced and repressed/uninduced conditions, respectively. Right: x-fold induction of the average fluorescence of 10,000 tracked events between induced and uninduced conditions.

For the comparative evaluation of the optogenetic switches, we focused on the following four criteria: (i) gradual inducibility (light dose response), (ii) basal expression in the corresponding OFF state, (iii) inducibility and dynamic range (fold-change between ON and OFF states), and (iv) heterogeneity of the population response determined at the single-cell level. As a reference, we first evaluated the UV-A-responsive P_tac_–LacI/NP-cIPTG system in *P. putida* (Figure 2A) whose basic functionality has already been demonstrated in this bacterium^15^. As expected, UV-A-mediated decaging of NP-cIPTG resulted in a strong induction of reporter gene expression. A more detailed characterization further revealed a gradual inducibility of the optochemical switch in dependence on the exposure time. Furthermore, flow cytometry data shows a very low basal expression (comparable to the WT control) and a high dynamic range of ∼270 fold (Figure 2H) but also a low expression heterogeneity within the population (Figure 2A). In addition, our results demonstrate that Tn7-based integration of the optogenetic expression cassette leads to a remarkably lower basal gene expression when compared to the plasmid-based system^15^.

The red-light sensitive OFF and ON switches showed a distinct response upon red-light illumination (λ = 625nm). For rR (OFF-switch, Figure 2B), increasing light intensities resulted in gradually decreased reporter fluorescence, with very low heterogeneity. On the other hand, for rD (ON-switch, Figure 2C), increasing light intensities led to rapid induction of reporter gene expression resulting in high mCherry fluorescence. Both mR and mD (Figure 2D,E) showed similar induction and repression profiles but exhibited opposite sensitivities at lower light intensity. In contrast to NP, red-light-responsive photoreceptors have slightly elevated basal activities in the OFF state, thereby gaining lower dynamic ranges (∼100 to ∼170 fold). Interestingly, the light-dose response (the light intensity needed for a certain level of expression), of these switches closely resemble the ones reported in *E. coli*.^3,4^ In contrast, the red-light switches reached up to 2.6-fold higher maximal fluorescence signals than the NP switch (Figure 2H), making them promising candidates for applications requiring high product yields.

The CcaS/R switch showed specific activation upon illumination at ∼525 nm, with strong repression at ∼625 nm. Since the GOI expression can be gradually induced across a wide range of light intensities, this optogenetic system enables a very precise and fine-tunable induction in *P. putida*. In addition, flow cytometry confirmed unimodal expression distributions, suggesting homogeneous population responses. Nonetheless, the total fluorescence, under fully induced conditions, is up to 16.6 times lower than for the red-light switches. In previous cases, the construct was delivered on a single vector, as for the original *E. coli* and adapted *P. putida* systems or as a dual vector design as for the later optimized versions for *E. coli*. Here, we chose genomic integration into the *att*Tn7 site, combining all genes in the corresponding optogenetic expression cassette that are necessary for the assembly of functional light switch (Figure 1A). This construction principle may minimize genetic instabilities and inconsistencies in expression behavior, but it also provides only one copy of the expression cassette, including the target gene. This could lead to reduced GOI expression levels. Recently, the de Lorenzo group successfully improved the CcaS/R system for optogenetic applications in *P. putida*. By optimizing regulatory sequences and balancing expression, an optimized CcaS/R cassette (pGreenL) could be generated that exhibited strongly increased reporter output and a higher dynamic range^12–14^.

The Dusk switch (Figure 2G) showed increasing, precisely adjustable repression of *mCherry* expression under blue light (λ = 465 nm) illumination, but the dynamic range was 2.8-fold lower compared to the NP and 1.3 up to 2.1-fold lower for the red-light-based switches (Figure 2H), mainly due to its slightly leaky expression at high light intensities. Furthermore, a slight expression heterogeneity could be observed upon blue light exposure (Figure 2 G). Nonetheless its dynamic range (∼100 fold induction) is still within reported ranges for *E. coli* and even five times higher than the reported dynamic range in *P. aeruginosa*^10,27^. Taking all results into account, we can conclude that all evaluated UV-A to RGB switches could successfully be implemented in *P. putida* and the performance reported in *E. coli* was generally preserved. Furthermore, each system exhibited individual strengths and weaknesses. NP can only be used as an irreversible switch but showed the lowest basal expression with the highest dynamic range in *P. putida.* The red-light-sensing, phytochrome-based switches showed the highest mCherry fluorescence signals in their ON-states, accompanied by homogenous expression behavior in all cells of the bacterial population. In contrast, the implementation of Dusk and the CcaS/R resulted in comparatively lower expression levels but demonstrated the best adjustability of the expression response. Overall, the established optogentic toolbox significantly expands the range of available regulatory parts for *P. putida* and enables new applications e.g., in synthetic biology where non-invasive control over gene expression with high spatio-temporal resolution is required.

### Multifactorial control and spatial patterning of gene expression

One of the most outstanding characteristics of optogenetics is its ability to control various biological functions *in vivo* with exceptional spatial and temporal resolution. To demonstrate the application potential of the optogenetic toolbox in *P. putida*, we first wanted to evaluate the spatially resolved expression of two different target genes by creating a bacteriograph, an image produced through patterned, light controlled reporter expression in a lawn of bacteria^37,38^(Figure 3).

**Figure 3.**
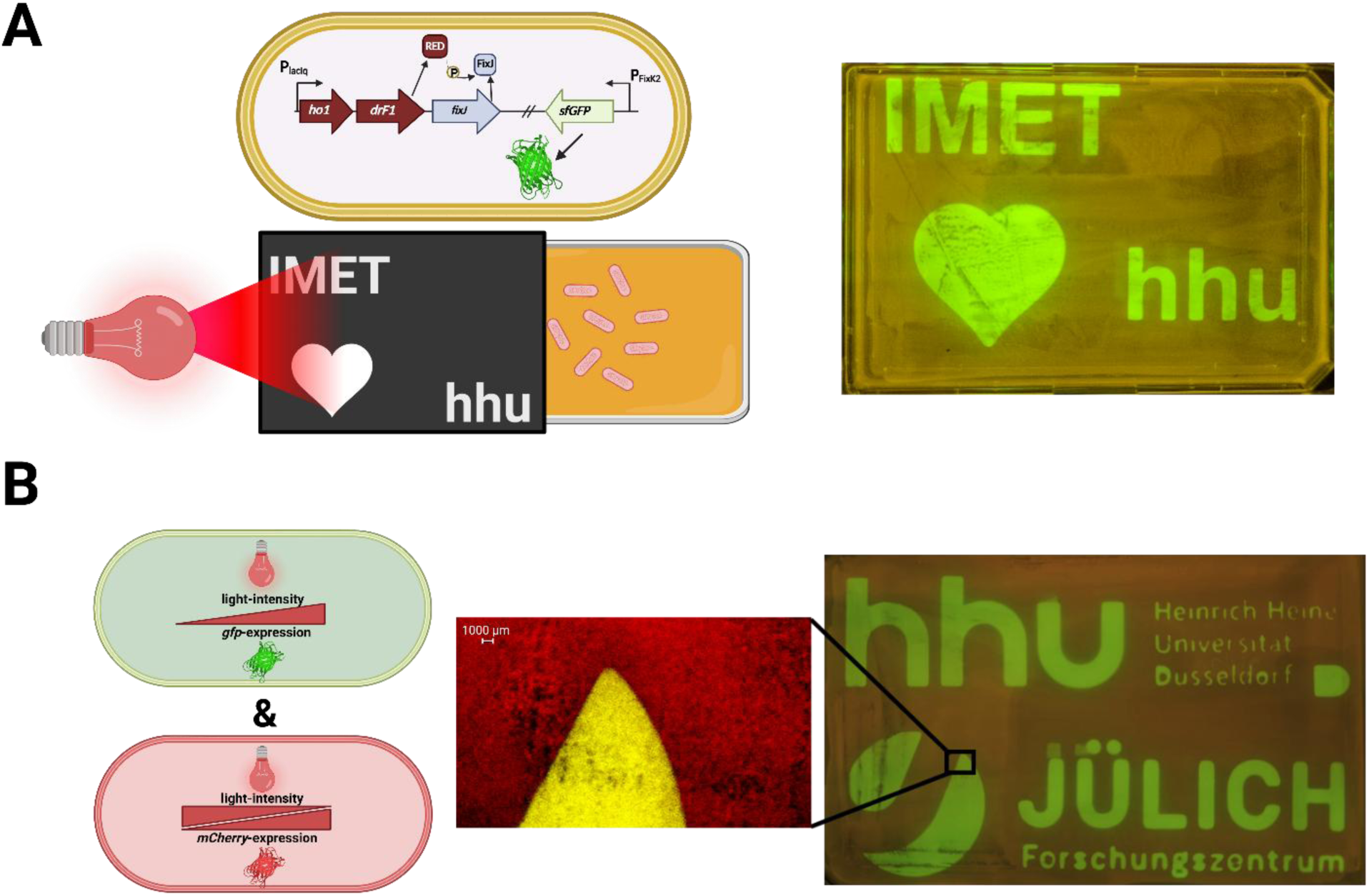
Single and multiplexed spatial patterning of gene expression using REDusk and rDERusk. (A, left side) Concept of experimental implementation of spatially structured fluorescence in a continuous bacterial layer induced by patterned illumination during cultivation. A digital light pattern was applied to agar plates that were confluently seeded with *P. putida* cells carrying the rDERusk switch controlling sfGFP expression. Partial illumination was used to gain locally restricted sfGFP expression patterns in the *P. putida* layer. Representative LB agar plates (overnight incubation under continuously patterned illumination) showed spatially confined sfGFP fluorescence corresponding to the projected pattern. (B) Multiplexed spatial control using *P. putida* harboring rDERusk (rD) controlling sfGFP expression and a second *P. putida* strain carrying the rREDusk (rR) switch controlling mCherry expression. Combined cultivation enables high spatiotemporal control of gene expression, with sfGFP fluorescence in illuminated and mCherry signals in non-illuminated regions.

*P. putida* carrying the *sfGFP* gene under the control of the red-light ON-switch rD was cultivated on an agar plate and was illuminated in defined patterns during cultivation. After incubation, only the illuminated areas showed sfGFP fluorescence (Figure 3 A). By co-cultivating the *P. putida* strain carrying the rD-sfGFP cassette with a *P. putida* strain carrying the complementary red light OFF-switch rR controlling mCherry gene expression, the differently illuminated areas showed clearly defined fluorescent sections within the confluent bacterial lawn. These results demonstrate that the optogenetic toolbox can be used to precisely control gene expression with high spatiotemporal resolution not only in liquid cultures but also in structured environments, thereby enabling new approaches for analyzing spatial pattering and dynamics of cellular functions as well as microbial interactions. As a potential extension, these optogenetic switches could be combined with microfluidic single-cell cultivation platforms, which previously have been used to quantify cellular responses to light induction, for example by activating photocaged IPTG^26,39^. Such systems enable high-resolution online analysis of biological heterogeneity at the single cell level ^40^ or the application of defined light gradients across the cultivation chip, allowing light-dose dependent kinetics to be studied^41^. Consequently, highly controlled monolayer microenvironments are suited to implement optogenetic switches in life cell imaging using time-lapse microscopy. Notably, the light intensities delivered to the cells during regular imaging are comparable to those needed to induce protein expression (Figure 2). Thus, imaging light itself could be exploited as a tuneable control signal, while also emphasizing the need for precise regulation of illumination conditions during microscopic studies, particularly with suitable optical filters and controlled light exposure.

### Establishing light-dependent control over pyoverdine biosynthesis in *P. putida*

So far, the new UV-RGB optogenetic systems were characterized in *P. putida* for the production of fluorescent proteins as reporters. While this allowed us to systematically compare performance features such as basal expression, inducibility, dynamic ranges and homogeneity, we next analyzed whether the established switches can also be applied to control more complex cellular processes. As a proof of concept, we chose the biosynthesis of pyoverdine (PVD), a secondary metabolite that plays an important role in iron acquisition in *P. putida.* Furthermore, PVD is of particular interest as it can mediate microbial and plant-microbe interactions, and has applications in bioremediation, pest control, and promoting plant growth^20,21,42–46^. The biosynthesis of PVD is naturally regulated by the iron availability (Figure S1). Under iron-limiting conditions, repression of the *pfrI* gene, which is mediated by the ferric uptake regulator Fur, is released. The formation of the alternative sigma factor PfrI results in the concerted expression of all PVD biosynthesis genes (*pvd*)^22^. To investigate, whether PVD production can be regulated by an optogenetic switch, we generated a *P. putida ΔpfrI* mutant strain, thereby abolishing the native Fur–PfrI dependent control of *pvd* gene expression. As reported previously, deletion of the *pfrI* gene resulted in a strongly impaired growth of *P. putida* under iron limited conditions (Figure S1 B,C)^47^. Based on these observations, we next tested whether the WT phenotype can be restored by light-induced complementation. To this end, the *pfrI* gene was reintroduced into the *ΔpfrI* mutant strain via the chromosomal integration of corresponding optogenetic expression cassettes (Figure 4).

**Figure 4.**
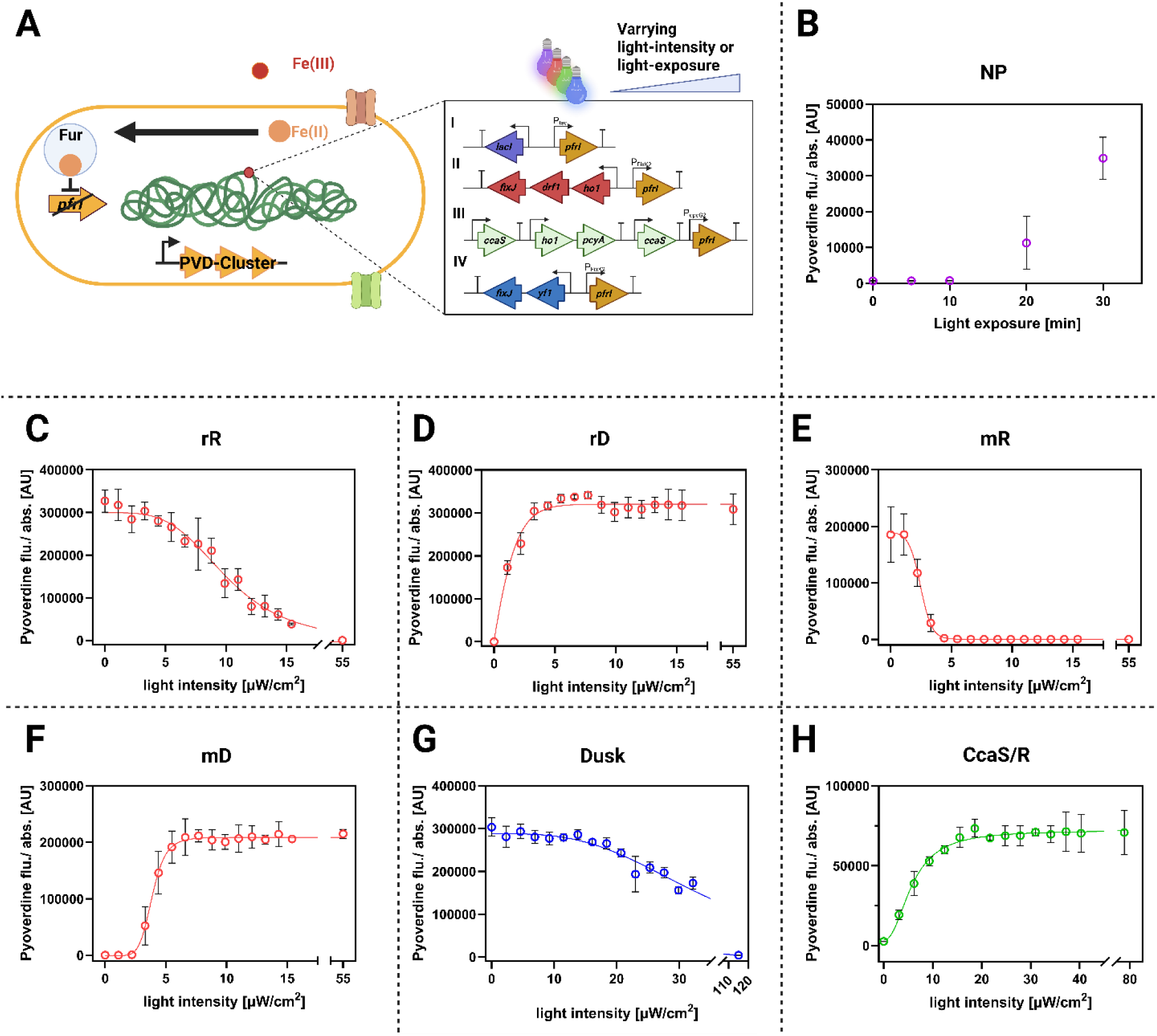
Establishing light-dependent control of pyoverdine biosynthesis in *P. putida*. (A) Light-dependent regulation of *pvd* gene expression in *P. putida*. In the WT strain the ferric uptake regulator (Fur) represses *pfrI* expression under iron-replete conditions, thereby blocking transcription of the *pvd* gene cluster. Under iron limitation, Fur repression is relieved and PfrI activates *pvd* gene expression (also see Figure S1). To decouple PVD production from iron availability, a *ΔpfrI* deletion mutant was generated and *pfrI* was reintroduced under the control of different optogenetic switches (I-IV), enabling iron-independent and light-mediated regulation of the native *pvd* gene cluster. (B-H) Quantification of PVD production in *P. putida ΔpfrI* strains carrying *pfrI* under the control of different optogenetic switches. PVD formation was analyzed by detecting the intrinsic fluorescence in the corresponding *P. putida* cultures (λ_ex_ = 398 nm, λ_em_ = 455 nm). Cultures were grown in 1 mL LB medium (30 °C, 1200 rpm) and illuminated with a programmable LED matrix (BioLight) at the indicated light intensities and wavelengths. The P_tac_-LacI system was illuminated after 4 hours for 0-30 mins with UV-A light (365nm ≈ 1mW/cm^2^). Data represents mean values ± SD from three biological replicates. (B) NP-cIPTG-based UV-A ON switch (∼365 nm), (C) red-light OFF switch rR (620–625 nm), (D) red-light ON switch rD (620–625 nm), (E) red-light OFF switch mR(620–625 nm), (F) red-light ON switch mD (620–625 nm), (G) green/red two-component system CcaS/R (∼522–525 nm and ∼620–625 nm), (H) blue-light OFF switch Dusk (465–467 nm).

To compare light-controlled PVD biosynthesis, Δ*pfrI* strains expressing *pfrI* under UV-A-, red-, green-, and blue-light control were analyzed across different illumination conditions (Figure 4B–H). The UV-A switch (P_tac_–LacI + NP-cIPTG) showed a very tight OFF state but unexpectedly low PVD production after induction, reaching only 0.6-fold of WT fluorescence under iron limitation (Supplementary Figure 1, Figure 4B). This reduced siderophore yield may result from UV exposure, which has been reported to reduce pyoverdine production^48^. In contrast, red-light systems provided the strongest and most controllable responses. The red-light ON switches rD and mD yielded the highest PVD production upon illumination, reaching 5.6-fold and 3.9-fold higher fluorescence than the iron-limited WT, while maintaining low basal activity in the dark (Figure 4D,F). Conversely, the OFF switches rR and mR strongly repressed PVD synthesis under light and promoted high production in the dark, achieving 6.2-fold and 3.4-fold higher fluorescence than the WT (Figure 4C,E). As observed previously, the r and m variants differed in their light sensitivity, again highlighting the importance of switch selection for specific applications. The CcaS/R system enabled reliable green-light activation and red-light repression, although total PVD production remained moderate (1.2-fold above WT; Figure 4G). In contrast, the blue-light OFF switch Dusk showed weaker repression at intermediate light intensities, suggesting that higher light intensities are required for precise tuning of PVD biosynthesis (Figure 4H). While iron-independent constitutive PVD production was previously achieved in *P. putida* by the constitutive expression of *pfrI*^22^, and inducible PVD production through bypassing the control through Fur was demonstrated in *P. aeruginosa*, by replacing the Fur-box upstream of the alternative sigma factor gene *pvdS* with an arabinose-inducible promoter^49^, here we established the first light-inducible *P. putida* strains for PVD biosynthesis - a new synthetic biology-based approach to achieve iron-independent, tunable, and color-specific control over PVD synthesis. Overall, these results clearly show that all optogenetic toolbox systems can be used to implement light-control over biosynthetic pathways in *P. putida*.

After establishing multicolor optogenetic expression modules in *P. putida* and light-mediated control over PVD production, we next wanted to elucidate whether this approach could also be used to achieve temporal control of PVD-based, microbial interactions under iron-limiting conditions. To this end, we focused on the two best-performing red-light switches, rR (OFF) and rD (ON), and investigated their ability to dynamically modulate siderophore production in liquid cultures as well as in agar plate-based interaction assays (Figure 5).

**Figure 5.**
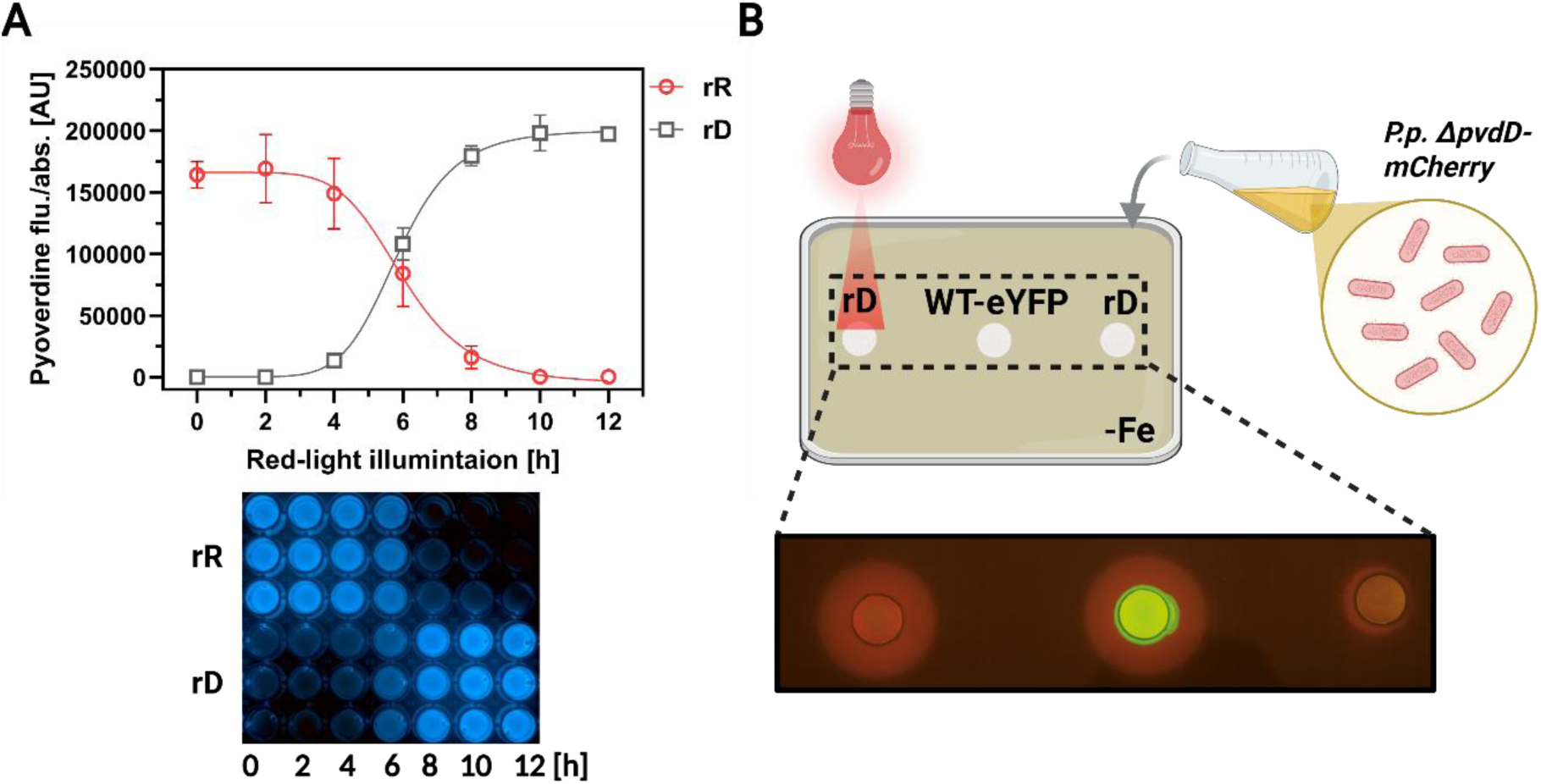
Light-dependent pyoverdine production enables spatial and temporal control over microbial interaction under iron limitation. (A) Light dose-response analysis of PVD biosynthesis in *P. putida ΔpfrI* expressing *pfrI* under control of the red-light OFF switch rREDusk (rR, red symbols) compared the red light ON switch rDERusk (rD, grey symbols). Cultures were grown for 24 h in microtiter plates and were illuminated for different time periods (0 to 12 h) with red light (625nm, 55 [µW/cm^2^]. After cultivation, gradual PVD production was directly visible as blue fluorescence and was subsequently quantified (λ_ex_ = 398 nm, λ_em_ = 455 nm). Data represents mean values ± SD of three biological replicates. (B) An overnight cultivation of the PVD-deficient *P putida ΔpvdD* strain labeled with mCherry, was streaked onto iron depleted agar. In addition, whatman paper discs were placed on the plate, which were first soaked either with the PVD producing strain rDERusk (without fluorescent label) or the WT strain labeled with eYFP as a control. As indicated by the formation of a red-fluorescing halo surrounding the whatman discs, *P putida ΔpvdD* was able to grow under iron limitation, only in the presence of a PVD producing strain (i.e., the WT or the illuminated rDERusk strain).

In liquid culture, the two red-light switches show inverse regulation of PVD synthesis in dependence of the applied exposure time with an induction curve comparable to the light-intensity dependent induction profile: The rD strain (ON, grey symbols) exhibited low basal PVD fluorescence in the dark, a steep increase with increasing red-light exposure duration, and a plateau at longer illumination (> 10 h). In contrast, rR (OFF, red symbols) showed high PVD output in the dark (< 2, which is increasingly repressed under prolonged illumination. These results confirm that both ON and OFF switches can be applied by modulating the light intensity (Figure 4) or exposure time (Figure 5) for siderophore production in *P. putida*, offering complementary modes of regulation. Other switches, such as CcaS/R and Dusk and rR, also showed differences in PVD production depending on the duration of light exposure (Figure S2). The functional relevance of optogenetic PVD control was finally demonstrated in a microbial interaction assay. To this end, we used a *P. putida ΔpvdD* mutant that is unable to produce PVD and therefore can only grow on iron-depleted agar in the presence of exogenous supplied PVD. When co-cultured with rD under red-light illumination, the *ΔpvdD* strain labeled with mCherry showed robust growth, visible as a red fluorescent halo, confirming that light-driven PVD synthesis provides accessible siderophores for the PVD-deficient strain. In contrast, the unexposed rD strain failed to sufficiently support growth of the *ΔpvdD* mutant, demonstrating tight spatial control over PVD-based microbial interaction. As a positive control, the WT labeled with eYFP supports growth of *P. putida ΔpvdD* in the dark.

PVD is a public good in pseudomonads. It is secreted, diffuses, and can therefore be exploited by non-producers, thereby directly impacting cooperation, cheating and competition dynamics in iron-limited environments. In *P. putida*, these population defining effects were analyzed by co-cultivating a constitutively PVD producing strain with a PVD-deficient cheater strain, in order to analyze the impact of public good production, sharing and cheating.^22^ It could be shown, that PVD producers initially have an advantage over cheaters during growth, but that the producer population suffers from PVD cheating in a later growth phase. Comparable public-good studies are well established in *P. aeruginosa*, and showed that, under conditions where non-producers can freely exploit PVD of the producer, the producer population declines over time.^50–52^ Importantly, these studies also showed, that this interaction is highly context dependent, and can change due to factors such as, population composition, environmental structures and local nutrient availability. More recent work suggests, that the PVD investment is not always homogenous even among clonal cells, but can be temporally and spatially coordinated.^53,54^ In this context, the use of defined microfluidic environments offer a promising platform to investigate interactions between cells at the microscale ^55^. By enabling precise spatial and temporal control of culture conditions, these systems can help disentangle how local gradients, population structure, and environmental heterogeneity shape microbial interactions^56^. Combined with light-responsive regulation, they further allow targeted modulation of gene expression while supporting continuous monitoring of co-culture dynamics. Beyond microbial interaction studies, PVD has also emerged as a potential plant growth promoting agent.^45,57,58^ Besides just increasing iron availability for plants, PVD has been detected in root and shoot tissue, and has been reported to impact the root morphology of *Arabidopsis thaliana*, although it remains unclear when or by which mechanism these effects occur.^59–61^ These results highlighted the ecological relevance of PVD availability in bacteria-plant interactions. Optogenetic regulation of PVD production therefore provides a highly valuable tool to study spatial aspects and dynamics of these interactions.

### Transferring light-dependent control over pyoverdine-synthesis to *P. aeruginosa*

To test whether the new optogenetic expression cassettes can also be applied beyond the non-pathogenic chassis of *P. putida*, we finally transferred it to the clinically relevant human pathogen *Pseudomonas aeruginosa* PAO1. As mentioned above, *P. aeruginosa* is one of the best studied model organisms for PVD-mediated public good interactions and importantly PVD is also linked to virulence during mammalian infection processes^62–64^. To this end, we introduced the *pfrI* sigma factor gene from *P. putida* into PAO1 under the control of either the IPTG-inducible P_tac_–LacI system or the red-light responsive rR switch (Figure 6).

**Figure 6.**
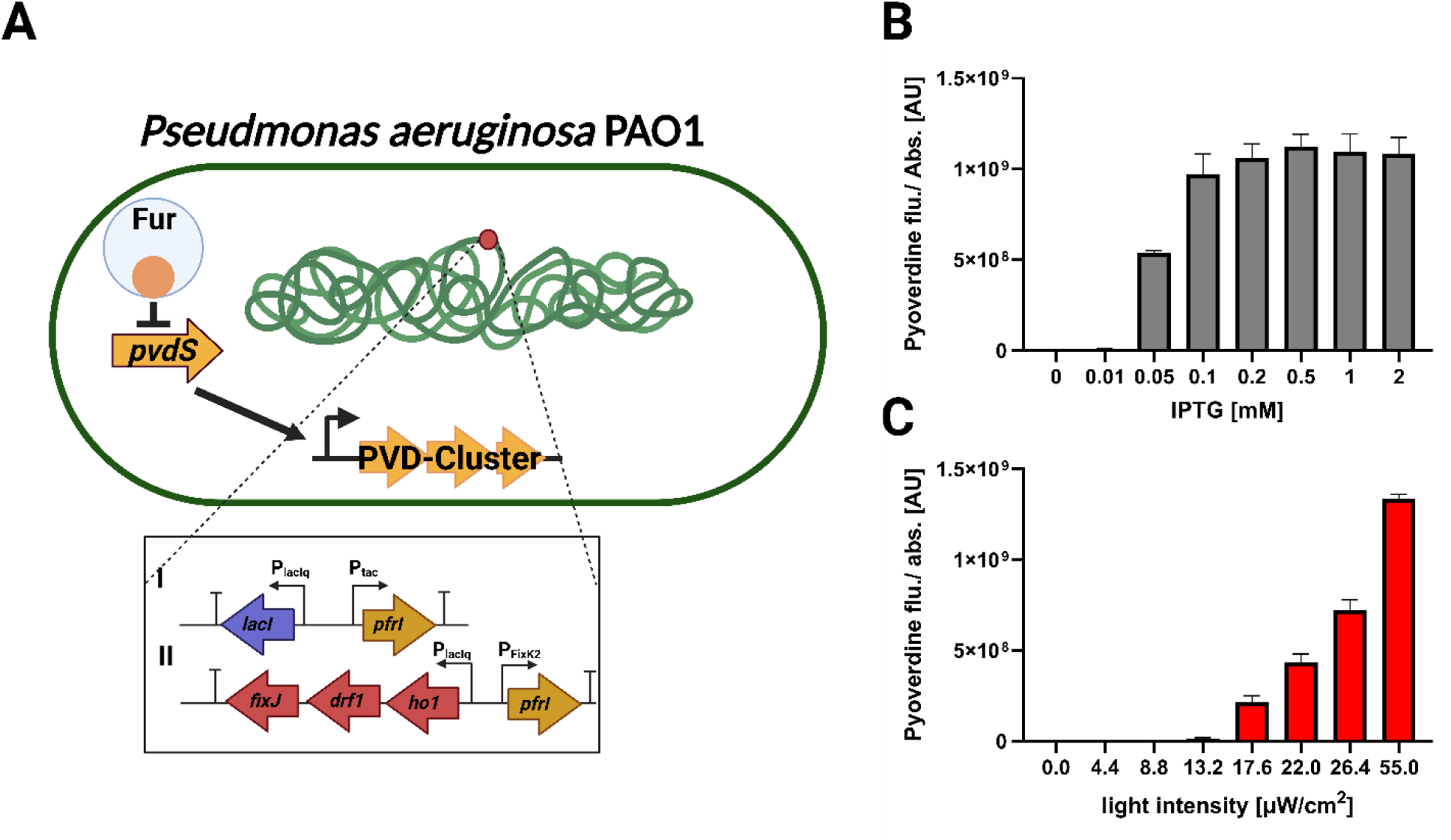
Optogenetic and chemical control of PVD production via heterologous expression of *pfrI* in *Pseudomonas aeruginosa* PAO1. (A) Schematic of PVD regulation in *P. aeruginosa*. Under iron limited conditions, Fur represses expression of the *pfrI* homologous gene *pvdS*. Similar to *P. putida*, the alternative sigma factor PvdS controls the expression of all *pvd* genes. To implement light-controlled PVD synthesis in the pathogenic model organism, two of the established optogenetic *pfrI* expression cassettes were integrated into the chromosome of *P. aeruginosa* via Tn7-madiated transposition. (I) IPTG-dependent and (II) red-light-mediated activation. (B) A *P. aeruginosa* strain carrying P_tac_–LacI–controlled *pfrI* was cultivated with increasing IPTG concentrations (0–2 mM), resulting in gradually elevated PVD production. (C) A strain harboring the REDusk (rD) system for controlling *pfrI* expression was exposed to increasing red-light intensities, yielding light-intensity-dependent PVD production. PVD fluorescence (λ_ex_ = 398 nm, λ_em_ = 455 nm) was normalized to OD600. Data represent mean values ± SD (n = 3).

In *P. aeruginosa*, iron uptake and the biosynthesis of siderophores such as PVD are tightly regulated by the Fur–PvdS cascade. Because PfrI is homologous to PvdS, we reasoned that it might also activate the PVD gene cluster in *P. aeruginosa* PAO1. Indeed, the P_tac_–*pfrI* strain showed IPTG-dependent activation of PVD biosynthesis (Figure 7B), while the REDusk–*pfrI* strain enabled gradual, red-light-dependent induction of siderophore formation (Figure 7C), revealing that PfrI can trigger PVD production in *P. aeruginosa*. These data demonstrate that the optogenetic expression toolbox can easily be transferred to other *Pseudomonas* species and enables the implementation of light-controlled signal transduction cascades. Furthermore, the establishment of light- and inducer-dependent PVD regulation in *P. aeruginosa* can facilitate future studies investigating PVD related microbial interactions and other pathogenic processes with defined temporal and spatial control. Moreover, red light offers improved tissue penetration compared to shorter wavelengths, allowing non-invasive control of virulence-associated functions in host cells and tissues.^65^

## Conclusion

In this study, we developed an optogenetic toolbox for *Pseudomonas putida* that enables light-dependent regulation of gene expression using UV-, red-, green-, and blue-light sensitive switches. This toolbox expands the possibilities to investigate microbial functions and processes with high spatial and temporal precision. As a proof of concept, we demonstrated light-controlled biosynthesis of the siderophore PVD in *P. putida* and *P. aeruginosa*, a secondary metabolite with biotechnological, bioecological, and biomedical relevance. In combination with microfluidic cultivation platforms, these tools could enable precisely controlled, spatially resolved studies of light-regulated bacterial behavior and microbial interactions at the single-cell level. Together, these tools provide a versatile platform for synthetic biology applications in *Pseudomonas*, ranging from tunable secondary metabolite production to the study of microbial interactions and other light-regulated processes.

## Materials and Methods

### Culture conditions and strains

*Eschericia coli* strains PIR2 ^66^ (Invitrogen; Thermo Fisher Scientific, Waltham, MA, USA) and DH5α λpir ^67^ as well as recombinant PIR2 and DH5α λpir strains carrying the optogenetic plasmids containing an R6kγ origin of replication, were grown in 1 mL of Luria-Bertani (LB) medium ^68^ at 37 °C in FlowerPlates^®^ (Beckman Coulter, Life Sciences, Baesweiler, Germany) under constant shaking at 1200 rpm in an Eppendorf ThermoMixer^®^ C (Eppendorf SE, Hamburg, Germany). *Pseudomonas putida* KT2440 wild type ^69^ as well as strains carrying *att*Tn7-site insertions were grown in 1 mL of LB medium at 30 °C under the same shaking conditions. *Pseudomonas aeruginosa* PAO1 wildtype ^70^ as well as strains carrying an attTn7-site insertions were grown in 1 mL of LB medium at 37 °C under the same shaking conditions. Where appropriate, antibiotics were added for plasmid maintenance, selection of chromosomal attTn7-site insertions of expression cassettes, or counterselection of donor strains. For *E. coli,* kanamycin, gentamicin and ampicillin were used at final concentrations of 50, 10 and 100 μg/mL, respectively. For *P. putida* and *P. aeruginosa*, Irgasan, gentamicin, and kanamycin were used at final concentrations of 25 μg/mL each.

All bacterial strains used in this study and their genetic properties are listed in Table S3 (Supporting Information).

### Plasmid construction

Plasmids were constructed by In-Fusion cloning using the In-Fusion® HD Cloning Kit (Takara Bio, Kusatsu, Japan) according to the manufacturer’s instructions. DNA fragments were amplified by PCR using the primers listed in Supporting Information Table S4 and assembled into the respective PCR-amplified vector backbones. Assembly products were transformed into chemically competent *Escherichia coli* PIR2 cells by heat shock at 42 °C for 90 s. Transformants were recovered in LB for 24 hours at 37 °C and selected on 1% (w/v) LB agar containing the appropriate antibiotic.

Recombinant plasmids (pDusk, pREDusk, pDrDERusk, pDmREDusk, pDmDERusk) harboring the corresponding optogenetic expression cassettes and fluorescent reporter gene (from pmTn7_Ptac_mCherry) were first generated by inserting the mCherry gene downstream of the respective optogenetically controlled promoters (Figure S5). The resulting optogenetic reporter cassettes, comprising the corresponding optogenetic switch and its promoter-controlled *mCherry* reporter gene, were amplified by PCR and transferred into the mini-Tn7 delivery vector pBG^35^ For this purpose, the pBG backbone was PCR-amplified from pBG13, and the optogenetic reporter cassettes were inserted into its cargo region between the Tn7L and Tn7R ends. The resulting pBG-derived mini-Tn7 delivery plasmids (pmTn7Dusk-mCherry, pmTn7REDusk-mCherry, pmTn7DrDERusk-mCherry, pmTn7DmREDusk-mCherry, pmTn7DmDERusk-mCherry) contain the gentamicin- and kanamycin-resistance genes, an R6Kγ origin of replication, and the respective optogenetic reporter cassette flanked by the Tn7L and Tn7R ends. For multiplexed bacteriograph ^37,38^ experiments, mCherry in the construct carrying the rDERusk switch was replaced by sfGFP (Figure S7).

For optogenetic control of pyoverdine biosynthesis, the *pfrI* coding gene was amplified from genomic DNA of *P. putida* KT2440 and inserted downstream of the optogenetically controlled promoter by replacing *mCherry* (Figure S5). Constitutive fluorescent-labeling plasmids were derived from pBG13 in which *sfGFP*, expressed from the constitutive promoter P_Em7_ was replaced by either *mCherry* or *eYFP*. All newly assembled plasmids were verified by complete plasmid sequencing using Oxford Nanopore sequencing (Eurofins Genomics, Cologne, Germany) Detailed plasmid descriptions and primer sequences as well as schematic cloning strategies for the different constructs is provided in Supporting Information Tables S4 and supporting Figures S4 – S7 respectively.

### Generation and analysis of the *P. putida ΔpfrI* knockout mutant

The *P. putida* KT2440 *ΔpfrI* deletion mutant was generated using the I-SceI-mediated recombination and pQURE-assisted plasmid-curing workflow described previously^71^. The 573-bp upstream and 592-bp downstream homology regions, directly flanking the chromosomal *pfrI* gene, were amplified from genomic DNA of *P. putida* KT2440 using the primers listed in Supporting Information Table S4. The two homology regions were assembled into the suicide vector pSNW2^71^ in which the upstream and downstream regions were fused such that homologous recombination would result in complete deletion of the coding sequence of the *pfrI* gene. The resulting deletion plasmid pSNW2-ΔpfrI was propagated in *E. coli* DH5α λpir and transferred into *P. putida* KT2440 by electroporation. Primary co-integrants, resulting from the first homologous recombination event, were selected on LB-Agar plates containing 50 µg/mL kanamycin and identified by fluorescence conferred by the constitutively expressed GFP reporter encoded on the integrated pSNW2-ΔpfrI plasmid. To induce excision of the plasmid backbone by a second homologous recombination event, co-integrants were transformed with the I-SceI-expressing self-curing plasmid pQURE6^71^. I-SceI expression and subsequent plasmid curing were induced by cultivation in the presence of 2mM 3-methylbenzoate and 25 µg/mL gentamicin according to the published protocol^71^. Candidate colonies were screened by colony PCR using primers 44 and 45, which bind outside the deleted locus. Clones yielding the expected shortened *ΔpfrI* amplicon were selected, and the deletion was subsequently confirmed by Sanger sequencing of the corresponding PCR product. Confirmed clones were designated as *P. putida ΔpfrI*. To assess whether deletion of *pfrI* abolished pyoverdine (PVD) production under iron-limiting conditions, *P. putida* KT2440 wild type, *P. putida* Δ*pvdD* and *P. putida* Δ*pfrI* were cultivated in the presence or absence of the iron chelator 2,2′-bipyridyl which was used to reduce iron availability and induce PVD production in the wild type, as described previously^22,53^. Overnight precultures of *P. putida* KT2440 wild type and *ΔpfrI* mutant were grown in 1 mL LB medium in FlowerPlates® (Beckman Coulter, Life Sciences, Baesweiler, Germany) at 30 °C and 1200 rpm. Precultures were used to inoculate 1 mL LB medium either without supplementation or supplemented with 0.25 mM 2,2′-bipyridyl (Sigma-Aldrich, Darmstadt, Germany) to an initial OD_600_ of 0.05. Cultures were incubated in FlowerPlates® in a BioLector I microbioreactor (Beckman Coulter Life Sciences, Baesweiler, Germany) for 24 h at 30 °C and 1200 rpm. After cultivation, 20 µL of each culture was transferred to a clear-bottom 96-well microtiter plate (Greiner Bio-One, Kremsmünster, Austria) and diluted with 80 µL phosphate-buffered saline (PBS) with a pH of 7.4. Absorbance at 600 nm as a measure of cell density and PVD-mediated fluorescence of the diluted samples were measured using an Infinite M1000 PRO plate reader (Tecan Austria GmbH, Grödig, Austria). PVD fluorescence was recorded using an excitation wavelength of 398 nm and an emission wavelength of 455 nm^22,72^.

### Genomic integration of optogenetic cassettes and fluorescent marker proteins

Chromosomal integration of the optogenetic expression cassettes and constitutive fluorescent-marker cassettes was performed by four-parental patch mating using a mini-Tn7-based integration system, as described previously^28,35,73^. Recipient strains were *P. putida* KT2440 wild type, *P. putida* Δ*pfrI*, *P. putida* Δ*pvdD*, and *Pseudomonas aeruginosa* PAO1. *E. coli* PIR2 carrying the respective pBG-derived mini-Tn7 delivery plasmid served as the cargo donor strain. These plasmids contained either an optogenetic expression cassette or a constitutively expressed fluorescent-marker cassette encoding mCherry or eYFP, positioned between the Tn7L and Tn7R ends. Two additional *E. coli* helper strains were included in the mating mixture. *E. coli* HB101 carrying pRK2013 supplied the conjugative transfer functions required for plasmid mobilization, whereas *E. coli* DH5α λpir carrying pTNS1 supplied the *tnsABCD* genes required for site-specific mini-Tn7 transposition into the chromosomal attTn7 site of the recipient strain^35^. Freshly streaked colonies of the recipient strain, the pBG-derived cargo donor strain, and both helper strains were combined on 1% (w/v) LB agar plates and incubated overnight at 30 °C. Following mating, cells were streaked onto 1% (w/v) LB agar containing gentamicin (25 μg/mL) and Irgasan (25 μg/mL) and incubated overnight at 30 °C. Gentamicin selected for recipient strains carrying the mini-Tn7 integration cassette, while Irgasan was used to counterselect the *E. coli* donor and helper strains. Individual gentamicin-resistant colonies were subsequently streaked onto 1% (w/v) LB agar containing gentamicin and Irgasan, as well as onto 1% (w/v) LB agar containing kanamycin (25 μg/mL) and Irgasan (25 μg/mL). Colonies exhibiting a gentamicin-resistant but kanamycin-sensitive phenotype were selected as candidate chromosomal mini-Tn7 integrants, because the gentamicin-resistance marker is transferred with the mini-Tn7 cassette, whereas the kanamycin-resistance marker remains on the non-integrated delivery-vector backbone. Correct insertion of the respective cassette at the chromosomal attTn7 site was confirmed using primers Tn7R and PglmS-down *via* colony PCR^28,73^.

### Creating a custom LED-matrix for cultivation in Flower Plates^®^

To enable individually programmable illumination of cultures in FlowerPlates® (Beckman Coulter Life Sciences, Baesweiler, Germany), a custom LED illumination device was constructed. The device consisted of 48 individually addressable WS2812B RGB LEDs arranged according to the well geometry of the FlowerPlate® and mounted in a custom-designed, 3D-printed housing to align one LED with each cultivation well and reduce optical interference between adjacent wells. The LEDs were controlled using an Arduino Uno microcontroller and the Adafruit NeoPixel library. Custom firmware was used to define the illumination wavelength, relative intensity and temporal illumination program independently for each well. During all experiments, LED brightness was restricted to the low-intensity settings required for optogenetic stimulation. The device layout, wiring scheme and list of materials are provided in Supporting Figure S3 and Supporting Table S1.

### Light-controlled cultivation and fluorescence measurements

For characterization of optogenetically controlled fluorescent reporter gene expression and pyoverdine production, recombinant *Pseudomonas* strains carrying the respective chromosomally integrated optogenetic reporter or effector cassettes (Table S3), were precultivated overnight under the corresponding non-inducing light conditions in 1 mL LB medium in FlowerPlates® (Beckman Coulter Life Sciences, Baesweiler, Germany). *P. putida* strains were precultivated at 30 °C, whereas *P. aeruginosa* strains were precultivated at 37 °C, with constant shaking at 1200 rpm in an Eppendorf ThermoMixer^®^ C (Eppendorf SE, Hamburg, Germany). For light-controlled cultivation, overnight cultures were inoculated into 1 mL LB medium in FlowerPlates® to an initial OD_600_ of 0.05. Cultures were incubated for 24 h at 30 °C for *P. putida* / 37 °C for *P. aeruginosa* and 1200 rpm using the custom LED matrix described above. Cultures were exposed to the inducing or non-inducing illumination programs specified in the corresponding figure legend. After cultivation, 20 µL of each culture was transferred to a clear-bottom 96-well microtiter plate (Greiner Bio-One, Kremsmünster, Austria) and diluted with 80 µL phosphate-buffered saline (PBS; pH 7.4). Cell densities were determined by measuring optical density at 600 nm. Measurements for *P. putida* cultures were performed using an Infinite M1000 PRO plate reader (Tecan Austria GmbH, Grödig, Austria), whereas measurements for *P. aeruginosa* cultures were performed using a SpectraMax iD3 plate reader (Molecular Devices GmbH, Munich, Germany). For strains carrying an mCherry reporter, fluorescence was measured using excitation and emission wavelengths of 575 and 610 nm, respectively. For experiments assessing pyoverdine production, PVD-associated fluorescence was measured using excitation and emission wavelengths of 398 and 455 nm, respectively, as previously described for PVD quantification in *P. putida* KT2440^22,72^. Fluorescence values were blank-corrected and normalized to OD_600_. Each condition was analyzed using three biological replicates.

### Flow-cytometric analysis of mCherry fluorescence

For single-cell analysis of optogenetic reporter gene expression, recombinant *P. putida* strains carrying a chromosomally integrated *mCherry* reporter gene under the control of an optogenetically regulated promoter were used (Supporting Information Table S3). Single colonies were used to inoculate overnight precultures in 1 mL LB medium in FlowerPlates® at 30 °C and 1200 rpm under the corresponding non-inducing light conditions. Precultures were subsequently used to inoculate 1 mL LB medium in FlowerPlates® to an initial OD_600_ of 0.05. Main cultures were incubated for 24 h at 30 °C and 1200 rpm using the custom LED matrix as described above and exposed to inducing or non-inducing illumination conditions. For flow-cytometric analysis, cultures were diluted in 1×PBS (pH 7.4) to an OD_600_ of 0.01. Samples were analyzed using an Amnis® CellStream™ flow cytometer (Cytek Biosciences, Fremont, CA, USA; formerly Luminex/Merck). For each sample, 10,000 events were recorded. Cell populations were gated based on forward scatter (FSC) and side scatter (SSC), followed by singlet selection. mCherry fluorescence was excited using the 561-nm laser and detected through a 611/31-nm bandpass filter. Data was analyzed using CellStream Analysis Software.

### Generation of fluorescent bacteriographs

Fluorescent bacteriographs^37,38^ were generated using fluorescent reporter expression controlled by DERusk and REDusk. Strains were precultivated overnight in 1 mL LB medium at 30 °C and 1200 rpm under non-inducing light conditions. Cultures were adjusted to an OD_600_ of 5 and 50 µL of the normalized cell suspension was spread evenly onto 1% (w/v) LB agar plates. Spatially restricted illumination was applied using a 3D-printed photomask containing the desired image patterns. The mask was positioned directly on the plate lid. Plates were illuminated with a wavelength of 625 nm and an intensity of 55 µW/cm^2^ for 24 hours at 30 °C. Non-illuminated areas of the plate served as internal non-induced controls. Following incubation, fluorescence was visualized using a FastGene Blue/Green LED Transilluminator (Nippon Genetics Europe GmbH, Düren, Germany), and images were acquired using a smartphone camera. Magnified images were acquired using a Nikon SMZ18 stereomicroscope equipped with NIS-Elements AR 5.3.

### Spatial pyoverdine-complementation assay

To investigate whether localized, light-induced restoration of PVD production could support growth of a PVD-deficient *P. putida* strain (Δ*pvdD*^73^) under iron-limiting conditions, a spatial complementation assay was performed. To this end, the eYFP-labelled wild-type control strain *P. putida eYFP* and the optogenetically controllable PVD production strain *P. putida ΔpfrI rD-pfrI* were used as PVD-providing strains. All strains were pre-cultivated overnight in 1 mL LB medium at 30 °C and 1200 rpm under non-inducing illumination conditions. Cultures were adjusted to an OD_600_ of 1. A 50 µL aliquot of the mCherry-labelled *P. putida* Δ*pvdD* culture was spread evenly onto a 1% (w/v) LB agar plate supplemented with 0.25 mM 2,2′-bipyridyl to impose iron limitation. Sterile Whatman® paper discs were dipped into the diluted culture of either the eYFP-labelled *P. putida* KT2440 culture or optogenetically controllable *P. putida ΔpfrI rD-pfrI* and placed at defined positions on the agar surface. The wild-type-loaded disc served as a PVD-providing control under iron limitation. Of the two discs loaded with *P. putida* Δ*pfrI*::attTn7-rDERusk-*pfrI*, only one disc was illuminated with 625 nm light at an intensity of with 55 µW/cm^2^ using the LED-matrix, whereas the second disc was covered with a mask and therefore maintained under darkness as a negative control. Plates were incubated for 24 h at 30 °C. Following incubation, mCherry- and eYFP-mediated fluorescence was visualized using a FastGene Blue/Green LED Transilluminator (Nippon Genetics Europe GmbH, Düren, Germany), and images were acquired using a smartphone camera.

## <ASSOCIATED CONTENT>

### Supporting Information

Plasmid maps, raw data, and CAD files for the BioLight Matrix are available from the corresponding author upon reasonable request. Plasmid maps and raw data will be deposited in a public repository upon submission of the manuscript for peer-reviewed publication.

## <AUTHOR INFORMATION> incl. authors contribution

S.P. designed, conducted, and evaluated the experiments; prepared the figures; drafted and wrote the manuscript. G.P. made major contributions to the design and construction of the BioLight Matrix. L.K. designed and conducted preliminary experiments and edited the manuscript. F.H., A.W., and T.H. conducted and evaluated preliminary experiments. M.P., L.W., M.B., D.K. provided feedback and edited the manuscript. T.D. acquired funding, supervised the study, designed experiments, and wrote the manuscript.

## <ACKNOWLEDGEMENTS>

The authors want to thank Aileen Krüger and Julia Frunzke (Institute of Bio- and Geosciences IBG-1: Biotechnology, Forschungszentrum Jülich GmbH, Jülich, Germany), for the helpful discussions and help with generating stereomicroscope images of bacteriograph experiments.

## <ABBREVIATIONS>

attTn7: attachment site of Tn7 transposon
AU: arbitrary units
cIPTG: photocaged isopropyl β-D-1-thiogalactopyranoside
DIP: 2,2′-dipyridyl
eYFP: enhanced yellow fluorescent protein
FSC: forward scatter
Fur: ferric uptake regulator
GOI: gene of interest
HK: histidine kinase
IPTG: isopropyl β-D-1-thiogalactopyranoside
LB: lysogeny broth
LOV: light-oxygen-voltage
mCherry: monomeric Cherry fluorescent protein
NP-cIPTG: 6-nitropiperonyl-caged IPTG
OD_600_: optical density at 600 nm
PfrI: alternative sigma factor regulating pyoverdine synthesis
PSM: photosensory module
PVD: pyoverdine
sfGFP: superfolder green fluorescent protein
SSC: side scatter
TCS: two-component system
WT: wild type

