## Supporting Information for "A multichromatic UV-RGB optogenetic toolbox for control of gene expression in *Pseudomonas putida*"

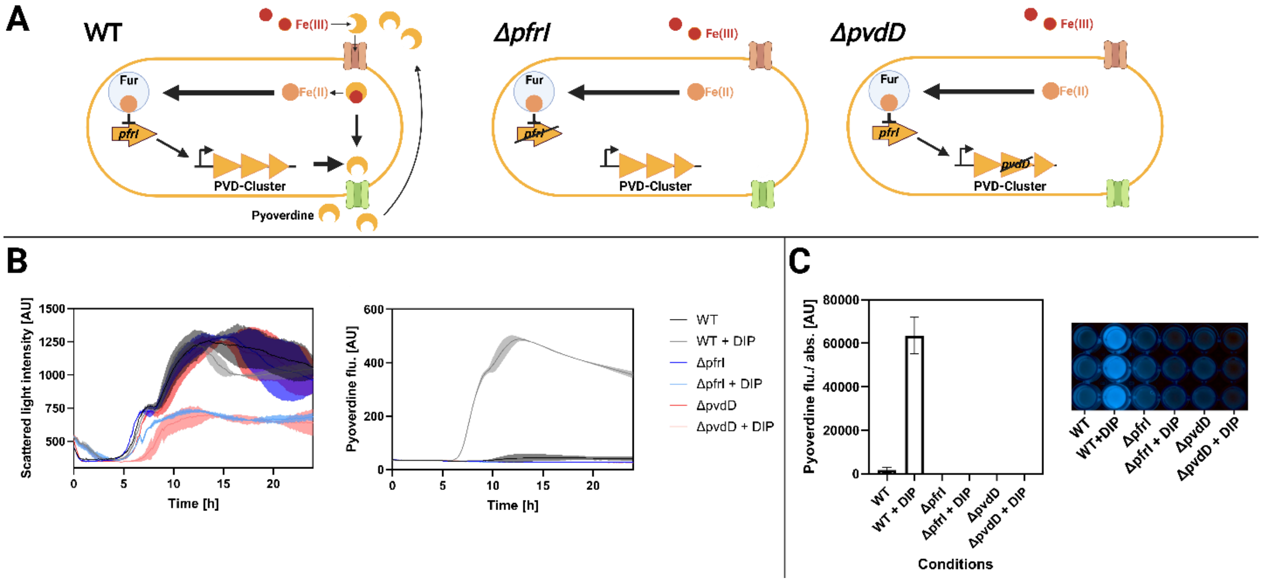


**Figure S1 Effect of *pfrI* and *pvdD* deletion on pyoverdine production and growth of *P. putida* under iron limitation.** (A) Schematic of pyoverdine (PVD) biosynthesis and iron acquisition in the wild type (WT) and the knockout strains Δ*pfrI* and Δ*pvdD*. In the *P. putida* WT strain, PfrI activates expression of PVD biosynthetic genes (incl. *pvdD*), enabling synthesis and secretion of the siderophore pyoverdine. The Δ*pfrI* mutant strain, lacking the alternative sigma factor, is not able to produce PVD. In the Δ*pvdD* strain the PVD biosynthesis is disrupted. (B, left) Growth of *P. putida* WT, Δ*pfrI*, and Δ*pvdD* strains measured as scattered light intensity over time under iron-sufficient (control) and iron-limited conditions. Iron limitation was imposed by adding 0.25 mM 2,2′-bipyridine (DIP). (B, right) Time course of pyoverdine fluorescence in culture supernatants ($\lambda_{ex}$ = 375 nm, $\lambda_{em}$ = 445 nm) measured online during growth. DIP strongly induces pyoverdine formation in the WT, whereas ΔpfrI and ΔpvdD are not able to synthesize the siderophore. (C) Quantification of PVD fluorescence normalized to cell density (mean ± SD) and qualitative PVD fluorescence detection of *P. putida* supernatants. The results confirm strong PVD synthesis only in the WT strain under iron limitation.

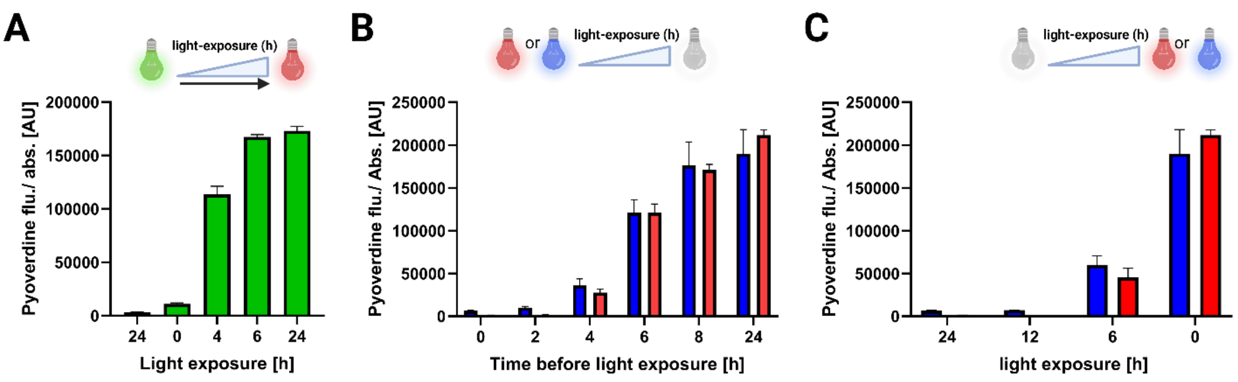


**Figure S2 Light-dependent regulation of pyoverdine production in *P. putida* under varying light exposure.** (A–C). PVD levels were quantified in culture supernatants by fluorescence ($\lambda_{ex}$ = 398 nm, $\lambda_{em}$ = 455 nm; values are shown as PVD fluorescence/cell density in AU). Bar plots summarize endpoint PVD formation after cultivation under the indicated exposure times, illustrating time--dependent control of PVD biosynthesis via optogenetic regulation of *pfrI* expression. Schematic icons above each panel indicate the applied light color and the timing of the light-to-dark (or light-to-light) transitions. Error bars represent mean values and SD of 3 biological replicates. (A) CcaS/R (green/red two-color switching). Cells were illuminated with green light for defined periods (0–24 h) to induce expression, then switched to red light for the remainder of cultivation. Longer green-light exposure results in gradually higher PVD accumulation, consistent with exposure-time dependent activation. (B) REDusk and Dusk systems (red- and blue-light illumination followed by darkness). Cultures were exposed to either red (REDusk) or blue (Dusk) light for different durations (0–24 h), then transferred to darkness for the rest of cultivation. Pyoverdine levels depend on the length of the initial light exposure and remain higher when illumination is shorter consistent with the OFF-switch properties of these switches. (C) Dark pre-incubation before light activation of REDusk and Dusk systems. Cultures were kept in the dark for varying times (0–24 h) prior to a light exposure (red or blue). Increasing the time spent in darkness before illumination gradually increases the induction.

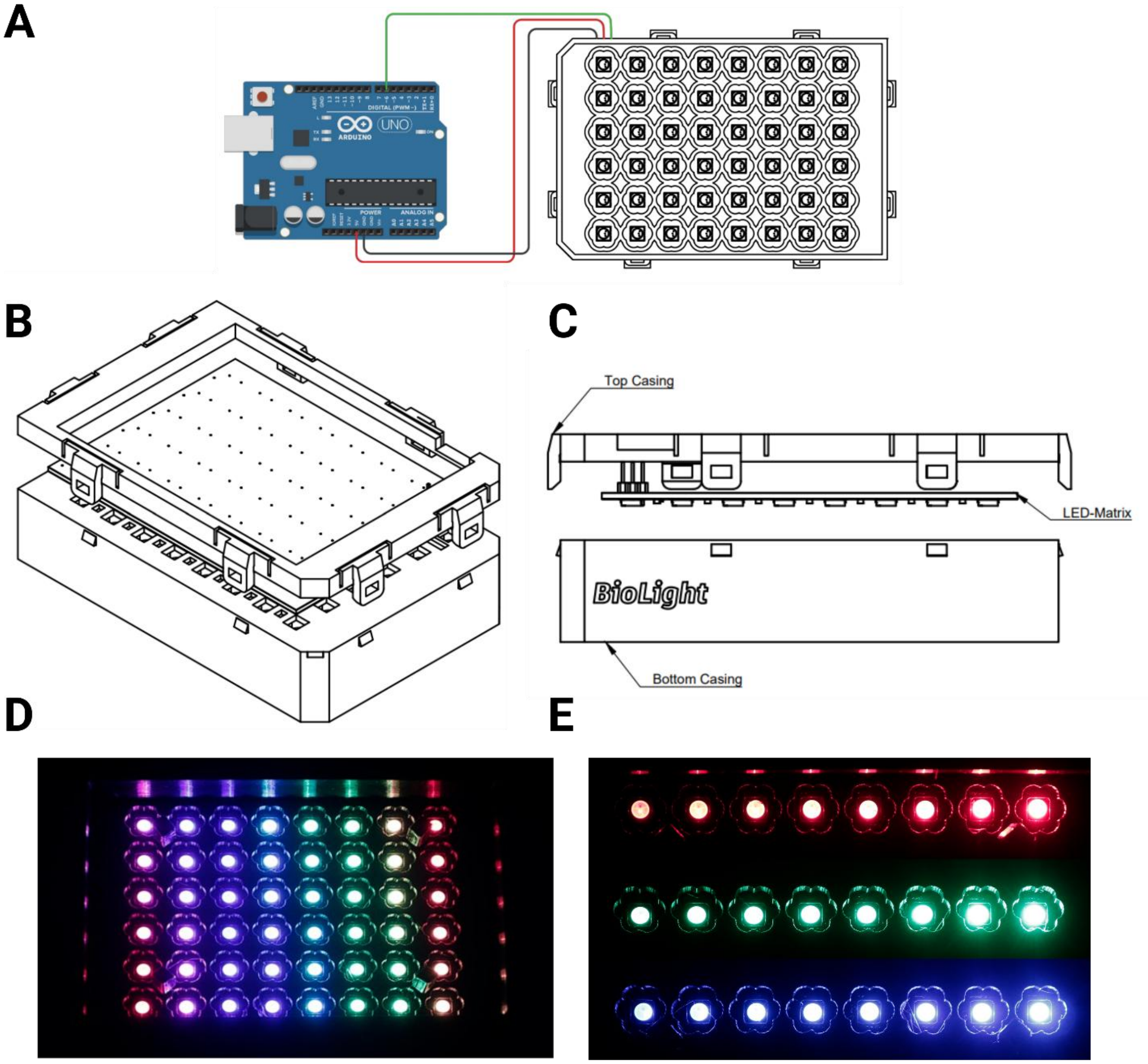


**Figure S3. Custom WS2812B LED–PCB matrix and housing for illumination of a FlowerPlate®.** A custom designed circuit board (PCB) carrying 48 individually addressable WS2812B RGB LEDs was designed such that each LED is aligned with a single well of a FlowerPlate®, enabling spatially defined, well-by-well light programming. The PCB is mounted in a 3D-printed casing that positions the LED array at a fixed distance above the plate and provides mechanical stability and light shielding. The LED matrix is controlled via a pre-programmed Arduino microcontroller. (A) Wiring overview of the Arduino-to-LED matrix connection (power, ground, and data line) used to drive the WS2812B array. (B) 3D view of the assembled illumination module showing the LED-PCB seated inside the 3D-printed enclosure designed to fit directly onto the FlowerPlate® format. (C) Side-view drawing of the housing, indicating top and bottom casing parts and the placement of the LED matrix within the enclosure to ensure consistent illumination geometry. (D) Photograph of the LED matrix operating a programmed “rainbow” test pattern, demonstrating individual addressability across the full 48-LED array. (E) Representative output of stepwise intensity modulation for each color channel (red, green, blue) showing graded light levels; peak emission wavelengths of the LEDs are Red 620–625 nm, Green 522–525 nm, and Blue 465–467 nm.

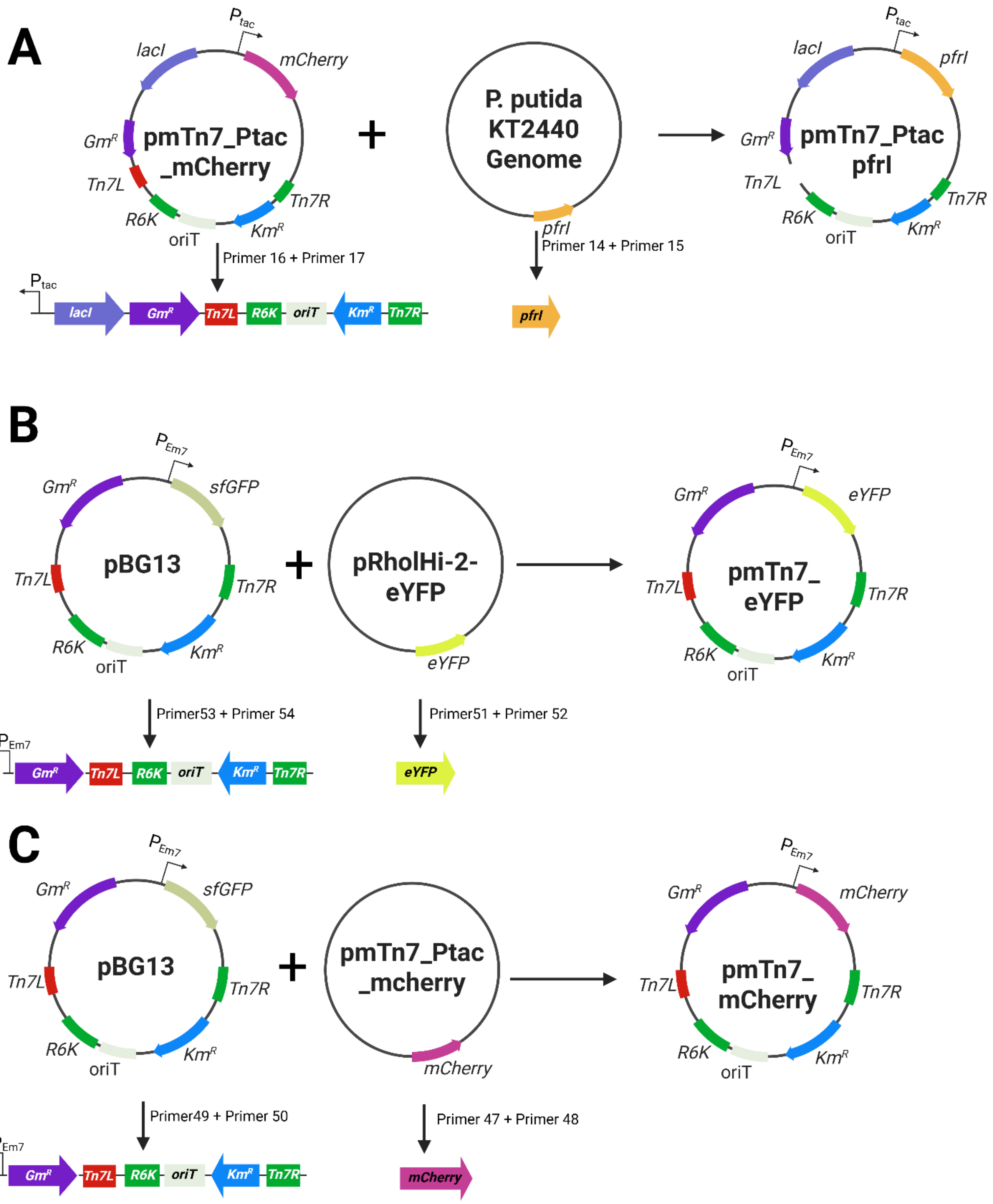


**Supplementary Figure S4. Construction of mini-Tn7 constitutive reporter expression and $P_{tac}$-pfrI expression vectors.** (A) Construction of plasmid pmTn7_Ptac_pfrI. The *pfrI* gene was obtained from the genome *of P. putida* KT2440 and introduced into the mini-Tn7 vector pmTn7_Ptac_mCherry, replacing the *mCherry* reporter gene. The resulting plasmid comprises *lacI*, a gentamicin resistance gene (*Gm*$^R$) the Tn7 left and right ends (Tn7L and Tn7R), the R6K origin of replication, an origin of transfer (oriT), a kanamycin resistance gene (*Km*$^R$) and *pfrI* under control of the $P_{tac}$ promoter. (B) Construction of plasmid pmTn7_eYFP. The *eYFP* reporter gene from pRholHi-2-eYFP was introduced into the mini-Tn7 backbone pBG13, resulting in plasmid pmTn7_eYFP. (C)

Construction of plasmid pmTn7_mCherry. The *mCherry* reporter gene from pmTn7_Ptac_mCherry was introduced into the mini-Tn7 backbone pBG13, resulting in plasmid pmTn7_mCherry. Genetic components are not drawn to scale.

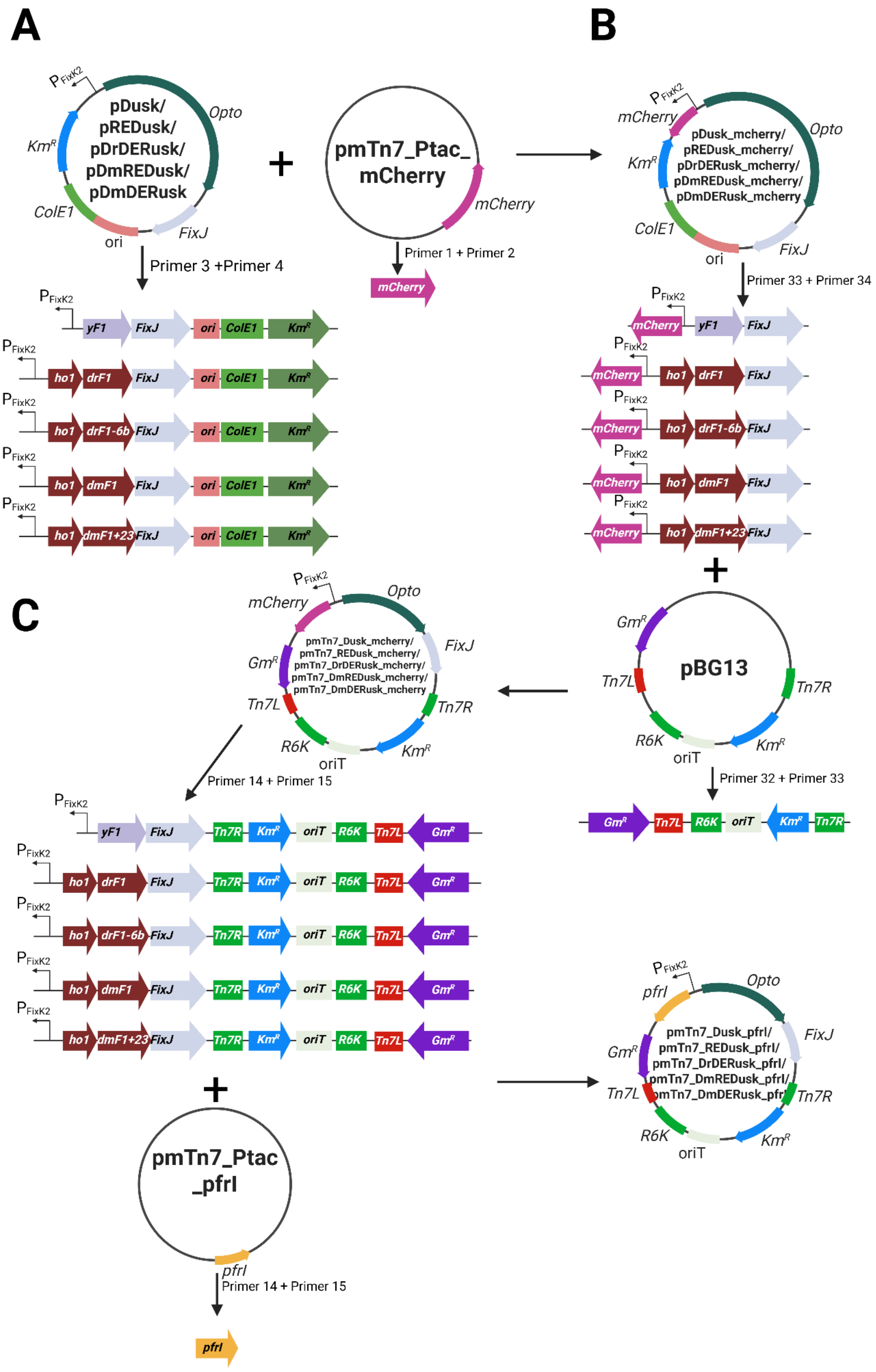
A
B
C
PFixK2
pDusk/
pREDusk/
pDrDERusk/
pDmREDusk/
pDmDERusk
Opto
KmR
ColE1
ori
FixJ
pmTn7_Ptac_
mCherry
mCherry
Primer 3 +Primer 4
Primer 1 + Primer 2
yF1
ho1
drF1
drF1-6b
dmF1
dmF1+23
pDusk_mcherry/
pREDusk_mcherry/
pDrDERusk_mcherry/
pDmREDusk_mcherry/
pDmDERusk_mcherry
Primer 33 + Primer 34
pBG13
GmR
Tn7L
Tn7R
R6K
oriT
Primer 32 + Primer 33
pmTn7_Dusk_mcherry/
pmTn7_REDusk_mcherry/
pmTn7_DrDERusk_mcherry/
pmTn7_DmREDusk_mcherry/
pmTn7_DmDERusk_mcherry
Primer 14 + Primer 15
pmTn7_Ptac
_pfrl
pfrl
pmTn7_Dusk_pfrl/
pmTn7_REDusk_pfrl/
pmTn7_DrDERusk_pfrl/
pmTn7_DmREDusk_pfrl/
pmTn7_DmDERusk_pfrl

**Supplementary Figure S5. Construction of mini-Tn7-based optogenetic reporter and expression vectors.** (A) Construction of the optogenetic *mCherry* reporter TCS vectors. The *mCherry* gene was isolated from plasmid pmTn7_Ptac_mCherry and introduced into the optogenetic expression cassette plasmid backbones carrying the blue-light-responsive YF1/FixJ module or red/far-red-light-responsive modules comprising *ho1* in combination with their respective PSM-histidinkinase fusion protein coding sequence (*drF1*, *drF1-6b*, *dmF1*, or *dmF1+23*) and *fixJ*. The resulting plasmids contain the respective optogenetic regulatory module upstream of *mCherry*, together with ColE1 origin of replication (ori), and a kanamycin resistance gene (*Km*$^{R}$). (B) Construction of the mini-Tn7 optogenetic *mCherry* TCS vectors. The optogenetic *mCherry* expression modules were transferred into the mini-Tn7 backbone pBG13, resulting in a set of plasmids containing the respective light-responsive regulatory module and *mCherry*. The resulting vectors additionally comprise a gentamicin resistance gene (Gm$^{R}$), the Tn7 left and right ends (Tn7L and Tn7R), the R6K origin of replication, an origin of transfer (oriT), and a kanamycin resistance gene (Km$^{R}$). (C) Construction of the mini-Tn7 optogenetic *pfrI* expression TCS vectors. The *pfrI* gene was obtained from pmTn7_Ptac_pfrI and introduced into the mini-Tn7 optogenetic vector set, replacing the *mCherry* reporter gene. The resulting plasmids contain *pfrI* under control of the respective blue- or red/far-red-light-responsive regulatory modules. Genetic components are not drawn to scale.

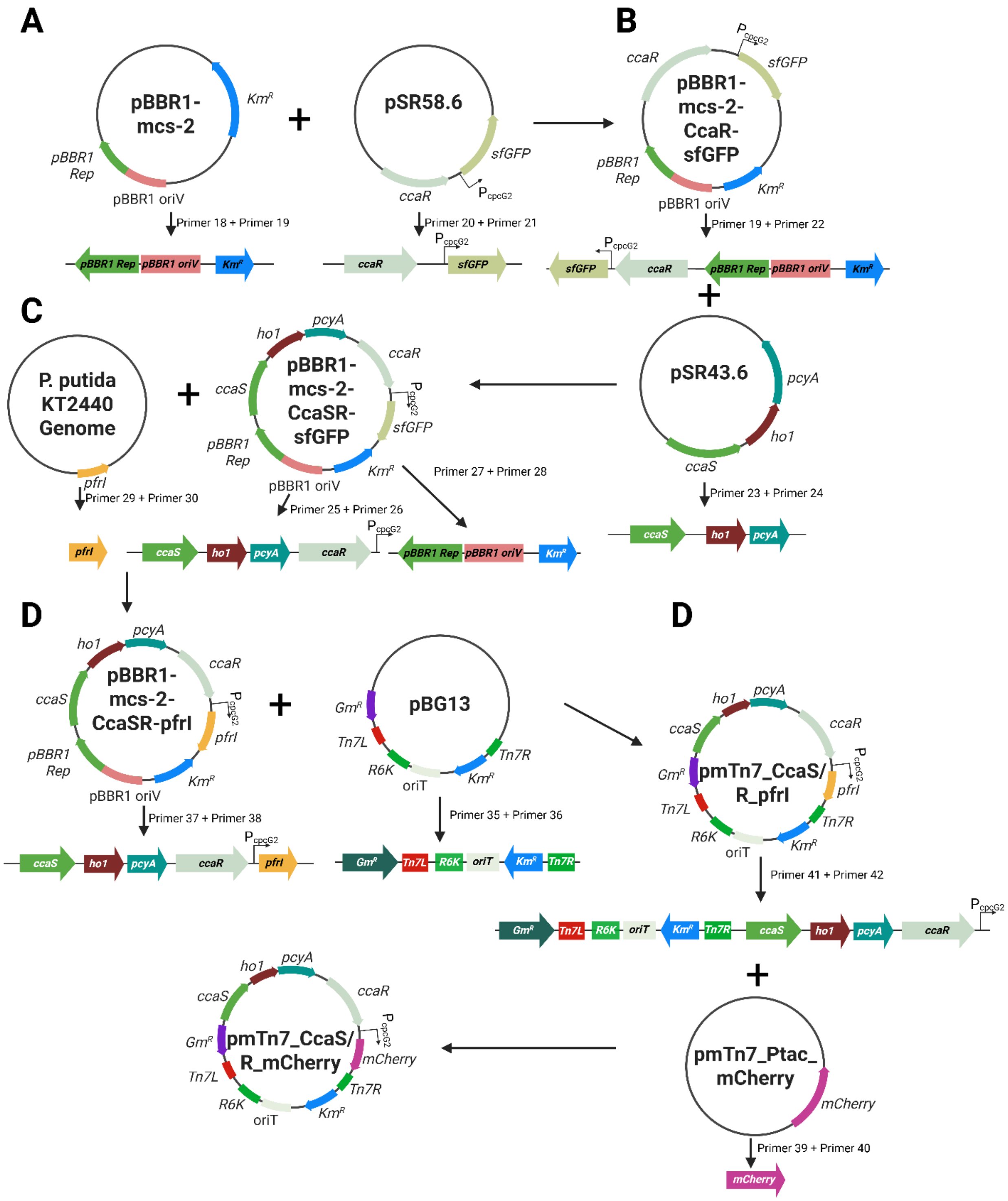


**Supplementary Figure S6. Construction of CcaS/CcaR-based green-light-responsive reporter and expression vectors.** (A) Construction of plasmid pBBR1-mcs-2-CcaR-sfGFP. The *ccaR-sfGFP* module was obtained from plasmid pSR58.6 and introduced into the broad-host-range vector pBBR1-mcs-2. The resulting plasmid contains the response regulator gene *ccaR* and the reporter gene *sfGFP*, together with the pBBR1 Rep region, pBBR1 oriV, and a kanamycin resistance gene ($Km^R$). (B) Construction of plasmid pBBR1-mcs-2-CcaSR-sfGFP. The sensory module comprising *ccaS*, *ho1*, and *pcyA* was isolated from plasmid pSR43.6 and introduced into pBBR1-mcs-2-CcaR-sfGFP, resulting in a complete CcaS/CcaR-based green-light-responsive *sfGFP* reporter vector. (C) Construction of plasmid pBBR1-mcs-2-CcaSR-pfrI. The *pfrI* gene was obtained from the genome of *P. putida* KT2440 and introduced into pBBR1-mcs-2-CcaSR-sfGFP, replacing the *sfGFP* reporter gene. The resulting plasmid contains *pfrI* under control of the CcaS/CcaR-responsive regulatory system. (D) Construction of the mini-Tn7 vector pmTn7_CcaSR_pfrI. The CcaS/CcaR-regulated *pfrI* expression module, comprising *ccaS*, *ho1*, *pcyA*, *ccaR*, and *pfrI*, was transferred into the mini-Tn7 backbone pBG13. The resulting vector additionally comprises a gentamicin resistance gene ($Gm^R$), the Tn7 left and right ends (Tn7L and Tn7R), the R6K origin of replication, an origin of transfer (oriT), and a kanamycin resistance gene ($Km^R$). (E) Construction of the mini-Tn7 reporter vector pmTn7_CcaSR_mCherry. The *mCherry* reporter gene was obtained from pmTn7_Ptac_mCherry and introduced into pmTn7_CcaSR_pfrI, replacing *pfrI*. The resulting plasmid enables CcaS/CcaR-dependent green-light-controlled expression of *mCherry*. Genetic components are not drawn to scale.

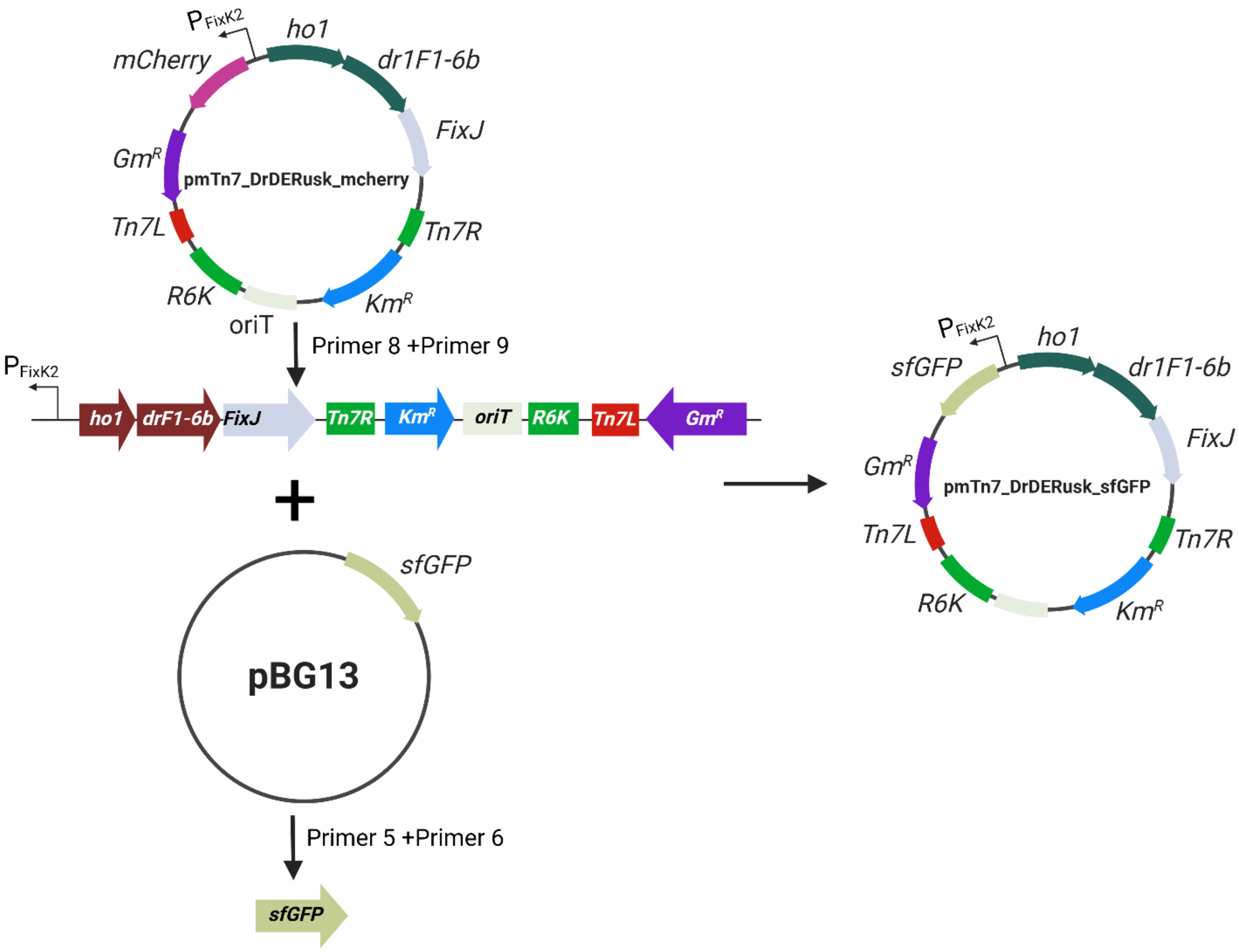


**Supplementary Figure S7 Construction of pmTn7_DrDERusk_sfGFP.** Construction of plasmid pmTn7_DrDERusk_sfGFP. The *sfGFP* reporter gene was obtained from pBG13 and introduced into pmTn7_DrDERusk_mCherry, replacing the *mCherry* reporter gene. The resulting plasmid contains a *sfGFP,* the optogenetic regulatory module comprising *ho1*, *drF1-6b*, and *FixJ*, together with a gentamicin resistance gene ($Gm^R$), the Tn7 left and right ends (Tn7L and Tn7R), the R6K origin of replication, an origin of transfer (oriT), and a kanamycin resistance gene ($Km^R$). Genetic components are not drawn to scale.

Table S1 List of components used for the LED Matrix and company of purchase

| Product | Company | Cat. number |
|---|---|---|
| LogiLink PA0139 USB Charger 32 W 6x USB-A | Conrad Electronic SE (Hirschau, Germany) | 1539271 - 62 |
| Arduino A000073 Board Uno Rev3 | Conrad Electronic SE (Hirschau, Germany) | 191789 - 62 |
| LogiLink USB-cable USB 2.0 USB-A | Conrad Electronic SE (Hirschau, Germany) | 973569 - 62 |
| Bambu Lab PLA Basic Black | IGO3D GmbH (Hannover, Germany) | PMBL-1001-005 |
| LED Matrix: | JiaLiChuang Co. Limited (JLCPCB), (Honkong, China) | Custom made |

Table S2 Used optogenetic switches

| System name | Light sensing module | Chromophore | On/off | System type | Reference |
| --- | --- | --- | --- | --- | --- |
| NP-IPTG | NP | NP | UV / Dark | Caged compound release | [1] |
| Dusk | YtvA | FMN | Blue /Dark | TCS | [2] |
| CcaS/R | CcaS | PCB | Green / Red & Dark | TCS | [3] |
| DrREDusk | DrBphP | Biliverdin | Far-Red & Dark/ Red | TCS | [4] |
| DrDERusk | DrBphP | Biliverdin | Dark/ Red & Far-Red | TCS | [5] |
| DmREDusk | DmBphP | Biliverdin | Far-Red & Dark/ Red | TCS | [5] |
| DmDERusk | DmBphP | Biliverdin | Dark/ Red & Far-Red | TCS | [5] |

Table S3 List of strains

| Strain | Description | Reference |
| --- | --- | --- |
| *E. coli* Pir2 | F - Δlac169 rpoS (Am) robA1 creC510 hsdR514 endA reacA1 uidA (ΔMlui)::pir | Life Technologies [6] |
| *E. coli* HB101 | F – mcrB mrr hsdS20(rB– mB– ) recA13 leuB6 ara-14 proA2 lacY1 galK2 xyl-5 mtl-1 rpsL20(SmR ) gln V44λ– | [7] |
| *E. coli* DH5α λpir | endA1 hsdR17 glnV44 (= supE44) thi-1 recA1 gyrA96 relA1 ϕ80dlacΔ(lacZ)M15 Δ(lacZYA-argF)U169 zdg-232::Tn10 uidA::pir+ | [8] |
| *P. putida ΔpvdD* | *P. putida* KT2440 with deletion of *pvdD* gene | [9] |
| *P. putida Δpfrl* | *P. putida* KT2440 with deletion of *pfrl* gene | This study |
| *P. putida* $P_{tac}$*-lacI-mCherry* | *P. putida* KT2440 with genomic attTn7 integration of: *Ptac-mCherry, LacI, GmR* | This study |
| *P. putida rR-mCherry* | *P. putida* KT2440 with genomic attTn7 integration of: *REDusk-mCherry, GmR* | This study |
| *P. putida mR-mCherry* | *P. putida* KT2440 with genomic attTn7 integration of: *mREDusk-mCherry, GmR* | This study |

| | | |
|---|---|---|
| *P. putida rD-mCherry* | *P. putida* KT2440 with genomic attTn7 integration of: *rDERusk-mCherry, GmR* | This study |
| *P. putida mD-mCherry* | *P. putida* KT2440 with genomic attTn7 integration of: *mDERusk-mCherry, GmR* | This study |
| *P. putida Dusk-mCherry* | *P. putida* KT2440 with genomic attTn7 integration of: *Dusk-mCherry, GmR* | This study |
| *P. putida CcaS/R-mCherry* | *P. putida* KT2440 with genomic attTn7 integration of: *CcaS/R-mCherry, GmR* | This study |
| *P. putida rD-sfGFP* | *P. putida* KT2440 with genomic attTn7 integration of: *rDERusk-sfGFP, GmR* | This study |
| *P. putida Δpfrl $P_{tac}$-lacI-pfrl* | *P. putida Δpfrl* with genomic attTn7 integration of: *Ptac-pfrl, LacI, GmR* | This study |
| *P. putida Δpfrl rR-pfrl* | *P. putida Δpfrl* with genomic attTn7 integration of: *REDusk-pfrl, GmR,* | This study |
| *P. putida Δpfrl mR- pfrl* | *P. putida Δpfrl* with genomic attTn7 integration of: *mREDusk-pfrl, GmR,,* | This study |
| *P. putida Δpfrl rD- pfrl* | *P. putida Δpfrl* with genomic attTn7 integration of: *rDERusk-pfrl, GmR,,* | This study |
| *P. putida Δpfrl mD- pfrl* | *P. putida Δpfrl* with genomic attTn7 integration of: *mDERusk-pfrl, GmR,* | This study |
| *P. putida Δpfrl Dusk- pfrl* | *P. putida Δpfrl* with genomic attTn7 integration of: *Dusk-pfrl, GmR,* | This study |
| *P. putida Δpfrl CcaS/R- pfrl* | *P. putida Δpfrl* with genomic attTn7 integration of: *CcaS/R-pfrl, GmR,,* | This study |
| *P. putida eYFP* | *P. putida KT2440* with genomic attTn7 integration of: *$P_{em7}$ eYFP, GmR* | This study |
| *P. putida ΔpvdD mCherry* | *P. putida KT2440* with genomic attTn7 integration of: *$P_{em7}$ mCherry, GmR* | This study |

Table S4 List of plasmids

| Plasmid | Description | Reference |
|---|---|---|
| pDrREDusk-MCS | Km$^R$, ColE1, *drF1*, *fixJ*, FixK2, MCS | [4] |
| DmREDusk-MCS | Km$^R$, ColE1, *dmF1*, *fixJ*, FixK2, MCS | [5] |
| DrDERusk-MCS | Km$^R$, ColE1, *drF1-6b*, *fixJ*, FixK2, MCS | [5] |
| DmDERusk-MCS | Km$^R$, ColE1, *dmF1+23*, *fixJ*, FixK2, MCS | [5] |
| pDusk | Km$^R$, ColE1, *yF1*, *fixJ*, FixK2 | [2] |
| pSR43.6 | Sm$^R$, *ho1, pcyA, ccaS* | [10] |
| pSR58.6 | Cm$^R$, *ccaR* | [10] |
| pRholHi-2-eYFP | pBBR1-MCS derivative, Km$^R$, Cm$^R$, $lacI_q$ $P_{tac}$, lacO, eyfp | [11] |
| pBBR1-mcs-2-CcaSR-sfGFP | Km$^R$, *ho1, pcyA, ccaS, ccaR, sfGFP, ori* (pBBR1) | This study |
| pDrREDusk_mCherry | Km$^R$, ColE1, *drF1*, *fixJ*, FixK2, *mCherry* | This study |
| pDmREDusk_mCherry | Km$^R$, ColE1, *dmF1*, *fixJ*, FixK2, *mCherry* | This study |
| pDrDERusk_mCherry | Km$^R$, ColE1, *drF1-6b*, *fixJ*, FixK2, *mCherry* | This study |
| pDmDERusk_mCherry | Km$^R$, ColE1, *dmF1+23*, *fixJ*, FixK2, *mCherry* | This study |
| pDusk_mCherry | Km$^R$, ColE1, *yF1*, *fixJ*, FixK2, *mCherry* | This study |
| pCcaS/R_pfrI | Km$^R$, *ho1, pcyA, ccaS, ccaR, pfrI, ori* (pBBR1) | This study |
| pBG13 | Km$^R$, Gm$^R$, *oriV* (R6K), pBG-derived, promoter $P_{em7}$, *msfGFP* | [12] |
| pmTn7_mCherry | Km$^R$, Gm$^R$, *oriV* (R6K), pBG-derived, promoter $P_{em7}$, *mCherry* | This Study |
| pmTn7_eYFP | Km$^R$, Gm$^R$, *oriV* (R6K), pBG-derived, promoter $P_{em7}$, *eYFP* | This study |
| pmTn7_DrREDusk_mCherry | Km$^R$, Gm$^R$, *oriV* (R6K), *drF1*, *fixJ*, FixK2, *mCherry* | This study |
| pmTn7_DmREDusk_mCherry | Km$^R$, Gm$^R$, *oriV* (R6K), *dmF1*, *fixJ*, FixK2, *mCherry* | This study |

| | | |
|---|---|---|
| pmTn7_DrDERusk_mCherry | Km$^R$, Gm$^R$, *oriV* (R6K), *drF1-6b*, *fixJ,* FixK2, *mCherry* | This study |
| pmTn7_DmDERusk_mCherry | Km$^R$, Gm$^R$, *oriV* (R6K), *dmF1+23*, *fixJ,* FixK2, *mCherry* | This study |
| pmTn7_pDusk_mCherry | Km$^R$, Gm$^R$, *oriV* (R6K), *yF1, fixJ,* FixK2, *mCherry* | This study |
| pmTn7_CcaS/R_mCherry | Km$^R$, Gm$^R$, *oriV* (R6K), *ho1, pcyA, ccaS, ccaR, mCherry* | This study |
| pmTn7_Ptac_mCherry | Km$^R$, Gm$^R$, *oriV* (R6K), $P_{tac}$, *lacI*, *mCherry, Tn7L, Tn7R* | [9] |
| pmTn7_DmDERusk_sfGFP | Km$^R$, Gm$^R$, *oriV* (R6K), *dmF1+23*, *fixJ,* FixK2, *sfGFP* | This study |
| pmTn7_DrREDusk_pfrI | Km$^R$, Gm$^R$, *oriV* (R6K), *drF1*, *fixJ,* FixK2, *pfrI* | This study |
| pmTn7_DmREDusk_pfrI | Km$^R$, Gm$^R$, *oriV* (R6K), *dmF1*, *fixJ,* FixK2, *pfrI* | This study |
| pmTn7_DrDERusk_pfrI | Km$^R$, Gm$^R$, *oriV* (R6K), *drF1-6b*, *fixJ,* FixK2, *pfrI* | This study |
| pmTn7_DmDERusk_pfri | Km$^R$, Gm$^R$, *oriV* (R6K), *dmF1+23*, *fixJ,* FixK2, *pfrI* | This study |
| pmTn7_pDusk_pfrI | Km$^R$, Gm$^R$, *oriV* (R6K), *yF1, fixJ,* FixK2, *pfrI* | This study |
| pmTn7_CcaS/R_pfrI | Km$^R$, Gm$^R$, *oriV* (R6K), *ho1, pcyA, ccaS, ccaR, pfrI* | This study |
| pmTn7_Ptac_pfrI | Km$^R$, Gm$^R$, *oriV* (R6K), $P_{tac}$, *lacI*, *pfrI, Tn7L, Tn7R* | This study |
| pQURE6·H | *oriV*(RK2), *XylS/Pm*→I-SceI and $P_{14g}$(*BCD2*)→*mRFP;* Gm$^R$ | [13] |
| pSNW2 | Suicide vector used for deletions in Gram-negative bacteria; *oriT, traJ*, *lacZα*, *oriV*(R6K), $P_{14g}$(*BCD2*)→*msfGFP*; *Km$^R$* | [13] |
| pSNW2_ΔpfrI | pSNW2, with flanking sequences for *pfrI* | This study |

Table S5 List of primers for plasmid construction

| Nr. | Primer name | Sequence 5'to3' | Usage |
|---|---|---|---|
| 1 | SP_160_IF_DuskSwapIns_fw | aggagatataccatgatggtgagcaagggc | PCR for mCherry insert with overhangs for In-fusion cloning into Dusk/REDusk and DERusk vectors |
| 2 | SP_161_IF_DuskSwapIns_rv | cctttcgggctttgtttacttgtacagctcgtcca | |
| 3 | SP_162_IF_DuskSwapBB_fw | gagctgtacaagtaaacaaagcccgaaagg | PCR for backbone with overhangs for In-fusion cloning with mCherry insert for Dusk/REDusk and DERusk vectors |
| 4 | SP_163_IF_DuskSwapBB_rv | gcccttgctcaccatcatggtatatctccttcttaaagt | |
| 5 | SP_268_IF_GFPinOP_ins_fw | cctttcgggctttgtttatttgtagagttcatccatg | PCR for sfGFP insert with overhangs for In-fusion cloning into Dusk/REDusk and DERusk vectors |
| 6 | SP_269_IF_GFPinOP_ins_rv | aggagatataccatgatgatcatgggaattcataa | |
| 8 | SP_270_IF_GFPinOP_BB_fw | aattcccatgatcatcatggtatatctccttcttaaag | PCR for backbone with overhangs for In-fusion cloning with sfGFP insert for Dusk/REDusk and DERusk vectors |
| 9 | SP_271_IF_GFPinOP_BB_rv | gaactctacaaataaacaaagcccgaaagg | |
| 10 | SP_276_IF_Pf_in_op_Ins_fw | cctttcgggctttgttcaggcctggcgact | PCR for pfrI insert with overhangs for In-fusion cloning into Dusk/REDusk and DERusk vectors |
| 11 | SP_277_IF_Pf_in_op_Ins_rv | aggagatataccatgatggcggaacaactatc | |
| 12 | SP_278_IF_Pf_in_op_BB_fw | tagttgttccgccatcatggtatatctccttcttaaagt | PCR for backbone with overhangs for In-fusion cloning with pfrI insert for Dusk/REDusk and DERusk vectors |
| 13 | SP_279_IF_Pf_in_op_BB_rv | agtcgccaggcctgaacaaagcccgaaagg | |
| 14 | SP_40_IF_Pf_in_Pt_fw | caggaaacagaattcatggcggaacaactatc | PCR for pfrI insert with overhangs for In-Fusion cloning into pMtn7_Ptac_ backbone |
| 15 | SP_41_IF_Pf_in_Pt_rv | 'gaattttctagaacgtcaggcctggcgact | |

| | | | |
|---|---|---|---|
| 16 | SP_42_IF_Pf_BB_Pt_fw | agtcgccaggcctgacgttctagaaaattcgtcaa | PCR for pMtn7_Ptac backbone with overhangs for In-fusion cloning with pfrI Insert |
| 17 | SP_43_IF_Pf_BB_Pt_rv | tagttgttccgccatgaattctgtttcctgtgtga | |
| 18 | FH_1_pBBR_fw | ggtcatttcgaaccccagag | PCR for backbone with overhangs for In-fusion cloning with CcaR insert |
| 19 | FH_2_pBBR_rev | gcgttaatattttgttaaaattcgc | |
| 20 | FH_3_58.6_fw | acaaaatattaacgccattttcgccagatatcgacgtct | PCR for CcaR insert with overhangs for In-fusion cloning into pBTBX |
| 21 | FH_4_58.6_rev | gggttcgaaatgaccatccacgggtacctataaacgcag | |
| 22 | FH_5_pBBR_fw | cattttcgccagatatcgacg | PCR with FH_2_pBBR_rev for backbone with overhangs for In-fusion cloning with CcaS, Ho1 and pcyA insert |
| 23 | FH_6_43.6_fw | acaaaatattaacgctggagtctggcctcaaatacaaag | PCR for CcaS, Ho1 and pcyA insert with overhangs for In-fusion cloning into pBTBX-CcaR |
| 24 | FH_7_43.6_rev | tatctggcgaaaatgtccccgggtacctataaacgca | |
| 25 | SP_019_IF_ccasrBBP1_fw | cggctatttaacgaccctgccctgaaccgacgaccgggtc | PCR for three fragment In-Fusion cloning, to generate CcaS/R_pfrI |
| 26 | SP_020_IF_CcasrBBP1_RV | gtggatagttgttccgccatctagtatttctcctctttttaaaaatgcgatcc | |
| 27 | SP_021_IF_ccasrBBP2_fw | cggccagtcgccaggcctgaataataatctagaccaggcatcaaataaaacg | |
| 28 | SP_022_IF_CcasrBBP2_RV | gacccggtcgtcggttcagggcagggtcgttaaatagccg | |
| 29 | SP_023_IF_PFRI_CCASR_fw | aaaaagaggagaaatactagatggcggaacaactatccacaagtaagtgc | |
| 30 | SP_024_IF_PFRI_CCASR_RV | tgcctggtctagattattattcaggcctggcgactggccg | |
| 31 | SP_090_IF_miniTn7_rv | ttaattaagacgtcttgaca | PCR for miniTn7 backbone with overhangs for In-fusion cloning with Dusk/REDusk and DERusk |
| 32 | SP_091_IF_miniTn7_FW | attcgagctcggtacccggg | |

| | | | |
|---|---|---|---|
| 33 | SP_124_IF_pDau_R__mt7_FW | gacgtcttaattaacaaaaacccctcaagacccg | PCR for Dusk/REDusk and DERusk insert with overhangs for In-fusion cloning into miniTn7 Backbone |
| 34 | SP_125_IF_pDau_R__mt7_rv | gtaccgagctcgaatactttacgaaacacggaaac | |
| 35 | SP_132_IF_CCASR_Pfri_FW | cgtcttaattaatcagatttaacaaaaatttaacgcg | PCR for CcaS/R-pfrI insert with overhangs for In-fusion cloning into miniTn7 Backbone |
| 36 | SP_133_IF_CCASR_Pfri_RV | tagttgttccgccatctagtatttctcctctttttaaaaat | |
| 37 | SP_134_IF_CCASR_PfriBB_FW | atggcggaacaactatc | PCR for miniTn7 backbone with overhangs for In-fusion cloning with CcaS/R-pfrI |
| 38 | SP_135_IF_CCASR_PfriBB_RV | tgattaattaagacgtcttgaca | |
| 39 | SP_168_IF_Ccswap_Ins_fw | gaggagaaatactagatggtgagcaagggc | PCR for mCherry insert with overhangs for In-fusion cloning into CcaS/R-pfrI backbone |
| 40 | SP_169_IF_Ccswap_Ins_rv | gaattttctagaacgttacttgtacagctcgtcca | |
| 41 | SP_170_IF_Ccswap_BB_fw | gagctgtacaagtaacgttctagaaaattcgtcaa | PCR for CcaS/R-pfrI backbone with overhangs for In-fusion cloning with mCherry |
| 42 | SP_171_IF_Ccswap_BB_rv | gcccttgctcaccatctagtatttctcctctttttaaaaat | |
| 43 | AW_1_pfrI-dw-fw | cgacgcgtcgagatgactgcaagcaaac | PCR from genome 592bp downstream of pfrI for ligation cloning into pSNW2 vector |
| 44 | AW_2_pfrI-dw-rv | gctctagagcgcagcaccgcgaatacttg | |
| 45 | AW_3_pfrI-up-fw[14] | cggaattccgccgaagaacgccgccacgtaatctg | PCR from genome 573 bp up stream of pfrI for ligation cloning into pSNW2 vector |
| 46 | AW_4_pfrI-up-rv | cgacgcgtcgggaaatcaccttgtcacagagg | |
| 47 | SP_050_IF_mChFW | taaggaggttttctaatggtgagaagggcga | PCR for mCherry insert with overhangs for In-fusion cloning into pBG13 Backbone |
| 48 | SP_051_IF_mCh_rv | accgagctcgaattcttacttgtacagctcgtccatgc | |

| | | | |
|---|---|---|---|
| 49 | SP_052_IF_PBG14BB_fw | gagctgtacaagtaagaattcgagctcggtacccg | PCR for pBG13 backbone with overhangs for In-fusion cloning with mCherry insert |
| 50 | SP_053_IF_PBG14BB_rv | gcccttgctcaccattagaaaacctccttagcatgattaagatg | |
| 51 | SP_054_IF_eYFP_fw | taaggaggttttctaatggtgagcaagggcga | PCR for eYFP insert with overhangs for In-fusion cloning into pBG13 Backbone |
| 52 | SP_055_IF_eYFP_rv | accgagctcgaattccttacttgtacagctcgtccatg | |
| 53 | SP_056_IF_PBG14BB2_FW | agctgtacaagtaaggaattcgagctcggtacccg | PCR for pBG13 backbone with overhangs for In-fusion cloning with eYFP insert |
| 54 | SP_057_IF_PBG14BB2_rv | gcccttgctcaccattagaaaacctccttagcatgattaagatg | |

Table S6 Primers used for colony PCR to verify Tn7 integration

| Primer name | sequence | Reference |
| --- | --- | --- |
| Tn7R | CACAGCATAACTGGACTGATTTC | [14] |
| PglmS-down | GCACATCGGCGACGTGCTCTC | [14] |
